# A Capsid Tyrosine Residue Governs the Proteolytic Inactivation of Echovirus 11 in Lakewater

**DOI:** 10.64898/2026.02.16.706081

**Authors:** Josephine Meibom, Shotaro Torii, L Daniela Morales, Michael Zumstein, Tamar Kohn

**Author notes:** To whom correspondence should be addressed: Tamar Kohn.

## Abstract

Enteroviruses are environmentally transmissible human pathogens whose stability in natural waters varies widely, yet the molecular determinants underlying this variability remain largely unknown. Echovirus 11 (E11), a re-emerging cause of severe neonatal infections, is efficiently transmitted via contaminated water, making its environmental stability a critical factor in infection risk. Here we identify a single viral capsid residue that governs E11 susceptibility to inactivation by extracellular microbial proteases in freshwater. By combining virus decay measurements in lakewater with proteolytic-cleavage profiling, viral capsid structural analyses, and reverse genetics, we show that the presence of VP2.Y97 renders E11 highly sensitive to microbially-mediated proteolytic decay. Strikingly, this residue is absent from multiple enteroviruses with greater environmental stability, indicating that substitution at a single capsid position is sufficient to shift virus fate in natural waters. These findings reveal that fine-scale capsid architecture controls virus-microbe interactions in aquatic environments and establish a molecular mechanism linking capsid variation to environmental transmission potential among enteroviruses.

## Introduction

Echovirus 11 (E11), a member of the *Enterovirus betacoxsackie* (formerly *Enterovirus B*) species of the *Enterovirus* genus, is among the most prevalent enteroviruses detected in environmental^1^ and clinical settings^1–3^. While often causing mild symptoms in adults, fatal outbreaks of E11 in newborns have been recently reported^4–7^, emphasizing the current public health risk of this pathogen. Commonly transmitted via the fecal-oral route^8,9^, enteroviruses may be introduced into aquatic environments through sewage overflow and insufficiently or untreated wastewater discharge^8^. As such, these pathogens can spread to new individuals through contaminated surface waters^9^. The transmission of enteroviruses through contaminated water is, however, largely dependent on the capacity of the viruses to resist biotic and abiotic environmental stressors. While microbial communities present in aquatic ecosystems are known to play a key role in viral decay^10–13^, the structural characteristics that drive the susceptibility of viruses such as E11 to microbial inactivation in these environments is poorly understood.

Some freshwater microorganisms can cause viral inactivation through the secretion of proteases. These enzymes can cleave peptide bonds present in structural viral proteins and consequently compromise the integrity of the viral capsid^14–16^. While several proteases have been found to inactivate various members of different virus families^14–18^, two classes of microbial proteases, the metallo- and serine proteases, have been broadly associated with enterovirus decay^14,18^. Interestingly, the antiviral activity of microbial proteases appears to be virus type specific. For example, the serine protease subtilisin was found to inactivate coxsackievirus A9, but not coxsackievirus B5 (CVB5) or E11^16^. Similarly, neutral protease displayed antiviral activity against coxsackievirus A9 but not against poliovirus 1^14^. While proteases are thus central to the microbial-mediated decay of enteroviruses, the molecular basis for their virus type specificity remains unexplored.

Microbial communities in aquatic ecosystems secrete a diverse repertoire of proteases presenting varying antiviral activities^18^. Although several proteases that inactivate specific viruses have been characterized^15,19^, it remains challenging to fully identify the pool of active proteases present in a complex aquatic environment. Instead, we can characterize the functional profile of proteases using Multiplex Substrate Profiling by Mass Spectrometry (MSP-MS)^20^. This method uncovers the substrate specificity profile, or proteolytic fingerprint, of enzyme pools by identifying amino acid patterns surrounding peptide bonds that are preferentially cleaved. Such fingerprints enable general inferences about which protein sequences are susceptible to proteolytic decay, as demonstrated for the hydrolysis of antimicrobial peptides by wastewater proteases^21,22^. Extending this framework to viral proteins by relating proteolytic fingerprints to virus capsid sequence and structure could provide mechanistic insight into proteolytic viral decay.

Herein, we investigated the proteolytic inactivation of E11 in lakewater using an integrated biochemical, molecular, and structural approach. By linking the proteolytic fingerprint of a lakewater protease pool to capsid structural features, four surface-exposed residues of E11 proposed to be susceptible to proteolytic cleavage were identified, and their relevance for viral inactivation was tested using reverse genetics. We further placed these findings in a broader context of environmental virus fate by comparing the stability of E11 in lakewater to that of three structurally similar *Enterovirus betacoxsackie* types. Together, this work establishes how viral capsid architecture governs proteolytic virus inactivation in freshwater and provides a mechanistic basis for the substantial divergence in environmental stability among similar enterovirus types.

## Results

### Protease-mediated inactivation of E11 in lakewater

To evaluate the stability of E11 in lakewater, we incubated the virus in several Lake Geneva surface water samples taken over the course of 16 months and monitored infectivity over time (**Figure 1A & Supplementary Figure S1**). In all samples, we observed first-order inactivation, resulting in 1.2 to 4.5 log_10_ decay at 54 hours (inactivation rate constant *k* ranging from 0.05 ± 0.01 to 0.19 ± 0.01 h^-1^, **Supplementary Table S1**) in microbially active lakewater. Although inactivation in sterile lakewater was also observed - possibly due to antiviral chemical compounds^23–26^ present in lakewater or to microbial regrowth (**Figure 1A & Supplementary Figure S1**) – we consistently observed greater decay in microbially active lakewater across all samples. To verify the contribution of proteases to inactivation, we added the metalloprotease inhibitor GM6001 to two lakewater samples prior to the incubation of E11 (**Supplementary Figure S2**). In these samples, we observed a reduction in inactivation by up to 1 log_10_ unit, confirming the contribution of metalloproteases to E11 decay. These results corroborate the importance of microbial, and specifically proteolytic, mechanisms in the inactivation of E11 in lakewater.

**Figure 1.**
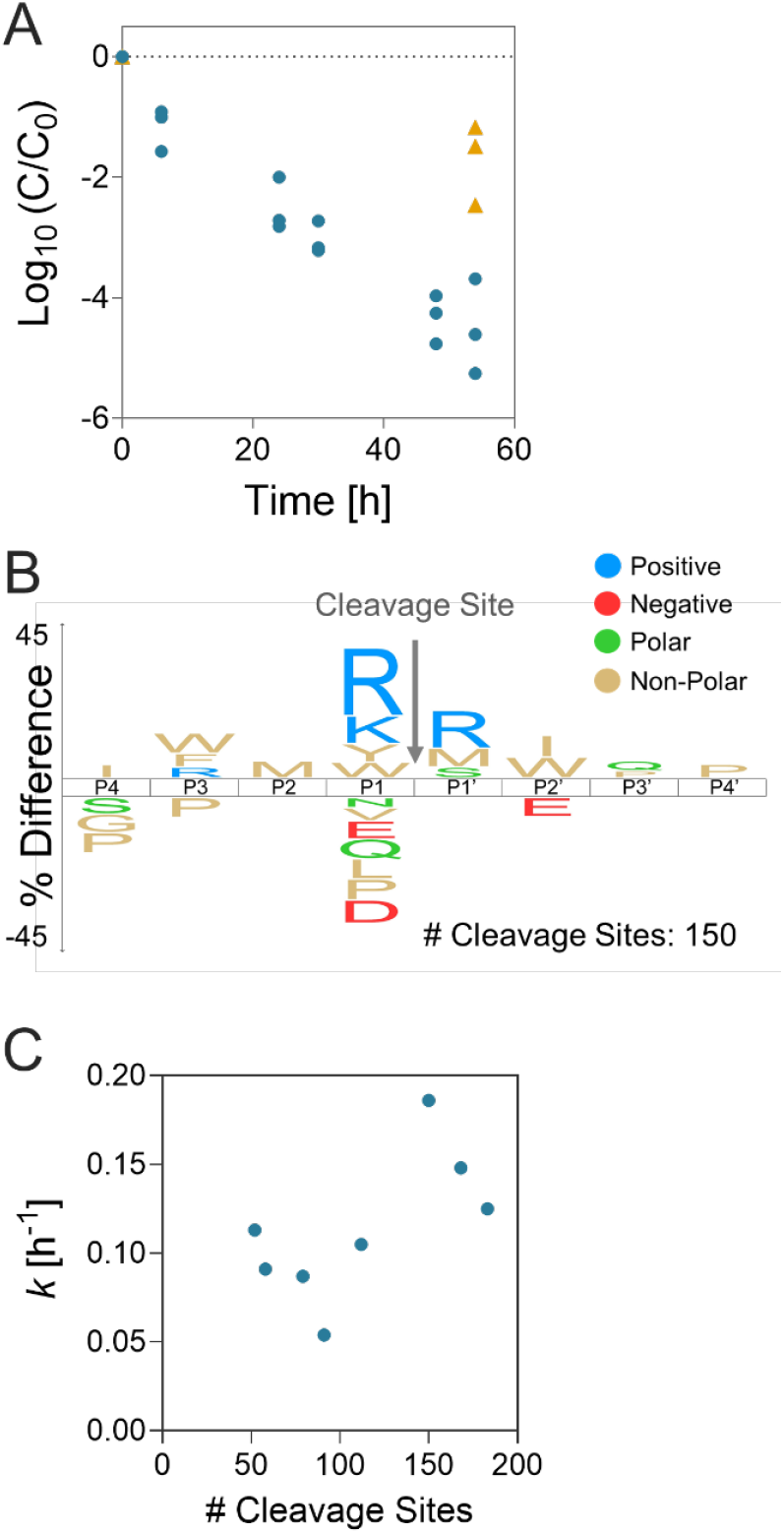
The proteolytic inactivation of E11 in lakewater **(A)** and the proteolytic fingerprint of the lakewater protease pool **(B).** Surface water was collected from Lake Geneva in September 2025. Individual data points of triplicate experiments are presented in (A). The number of proteolytic cleavage sites detected by the MSP-MS assay, reflective of the heterogeneity of the substrate specificity, is indicated in (B). **(C)** Correlation between the mean E11 inactivation rate constants *k* (**Supplementary Table S1**, n = 3) and the number of MSP-MS cleavage sites detected over 6 hours in each lakewater sample (Pearson correlation coefficient r = 0.625).

To characterize the substrate specificity of the microbial protease pool in Lake Geneva and to gain insight into the proteolytic inactivation of E11, we employed the MSP-MS assay. This method allows us to determine which amino acids are preferred or disfavored at the eight positions (P4-P4’) surrounding the cleavage site. We previously reported the proteolytic fingerprints of Lake Geneva collected between April 2024 and February 2025^27^. These fingerprints revealed a strong preference for cleavage when positively charged amino acids (arginine and lysine) and aromatic amino acids (tryptophan and tyrosine) were present N-terminal to the cleaved peptide bond (position P1). Conversely, cleavage was disfavored when proline, negatively charged (aspartic acid and glutamic acid), and some polar (threonine and glutamine) amino acids were present at P1. We here report consistent patterns in three additional lakewater samples collected in April 2025, July 2025, and September 2025 (**Figure 1B & Supplementary Figure S3**), indicating that the dominant features of the Lake Geneva proteolytic fingerprint are temporally conserved. Combined with the consistently observed E11 inactivation in lakewater, these findings suggest that the dominant proteolytic substrate specificities coincide with vulnerable sites on the virus capsid.

Despite the stability of the proteolytic fingerprints, we observed fluctuations in the number of distinct cleavage sites detected by the MSP-MS assay (**Supplementary Figure S3**), indicative of varying degrees of heterogeneity in the minor substrate specificities of the lakewater protease pool^27^. We found a positive correlation between the number of cleavage sites and the inactivation rate constant *k* (Pearson correlation coefficient r = 0.625, **Figure 1C**), suggesting that increased heterogeneity in the protease pool associates with E11 inactivation.

### Identifying potential proteolytic cleavage sites on the surface of E11

We used the above-determined proteolytic fingerprints to identify potential cleavage sites on the surface of E11 (**Supplementary Figure S4**). While the P1 residue is often crucial in determining the substrate specificity of proteases – as reflected by its dominance in the proteolytic fingerprints, **Supplementary Figure S3**) - other residues surrounding the cleaved peptide bond can also influence substrate recognition and cleavage^28^. Here, we selected all combinations of P1-P1’ (flanking the cleaved peptide bond) and P2-P1 (N-terminal to the cleaved peptide bond) amino acid pairs observed in the lakewater proteolytic fingerprints (**Supplementary Figure S3 & Supplementary Table S2**). We mapped these pairs, which capture the dominant substrate specificities of the lakewater protease pool, onto the viral protein sequences of E11. This mapping revealed a subset of P1 residues that were surface-exposed on the E11 capsid, as determined by analysis of the modeled virus ultra-structure (**Figure 2A**). We modeled the E11 capsid using the exact viral protein sequences of the laboratory E11 isolate used here, and the resulting structure closely aligned (**Supplementary Figure S5**, RMSD = 0.676 Å) with a previously resolved experimental E11 structure (PDB 6LA3^29^), supporting its use for this analysis.

**Figure 2.**
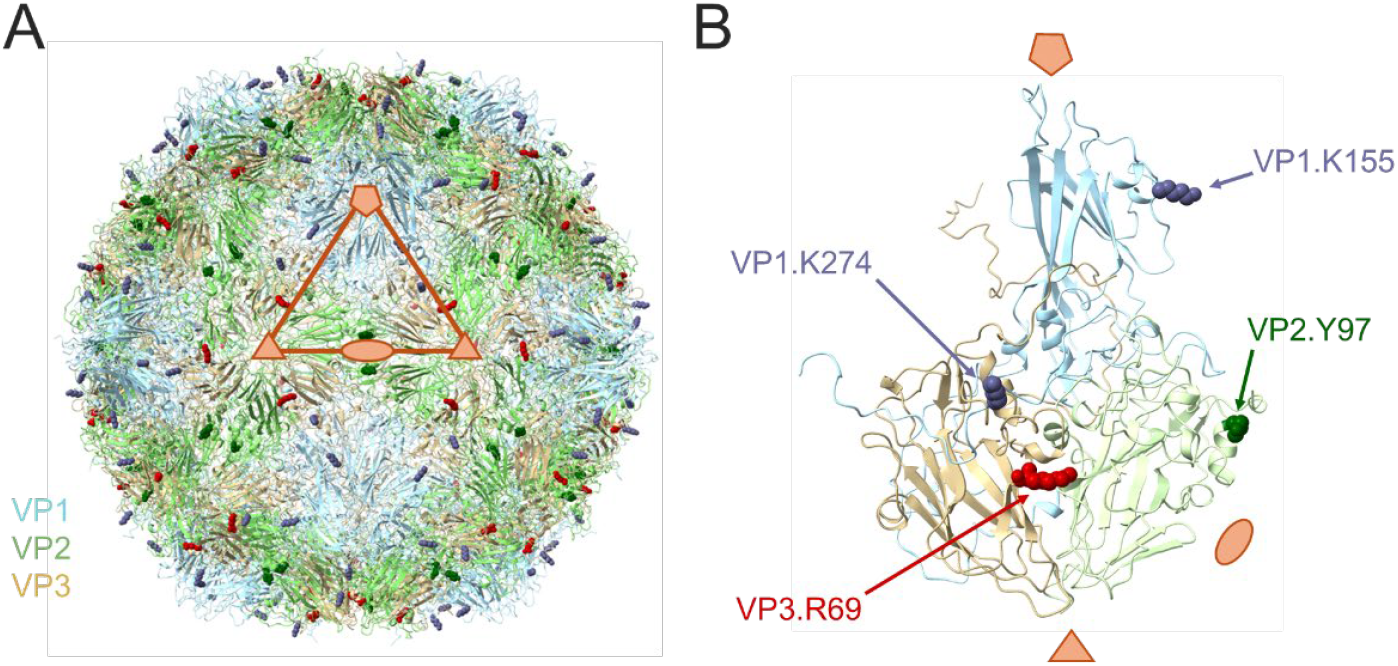
Selection of potential proteolytic cleavage sites on the surface of E11. Visual representation of **(A)** the entire E11 capsid and **(B)** an individual protomer highlighting the four potential proteolytic cleavage sites: VP1.K155 (K-S P1-P1’ match), VP1.K274 (K-R P1-P1’ match), VP2.Y97 (M-Y P2-P1 match), and VP3.R69 (R-I P1-P1’ and Y-R P2-P1 matches). The asymmetric unit (orange triangle in (A)) is indicated along with the 2-, 3-, and 5-fold icosahedral symmetry axes represented by an ellipse, triangle, and pentamer, respectively. Images were produced in ChimeraX^32^.

We subsequently designated P1 residues that were visually identified as fully surface-exposed as possible proteolytic cleavage sites. Based on this analysis, four residues were identified: VP1.K155, VP1.K274, VP2.Y97, and VP3.R69 (**Figure 2 & Table 1**). These residues were distributed across the external structural viral proteins (VP1-VP3) and were located on flexible, disordered regions (loops or the C-terminal sequence) suggesting that they are potentially accessible to proteases^16,30,31^.

**Table 1.** Selected amino acid residues for substitution and the corresponding E11 mutants produced based on the proteolytic cleavage site selection process detailed in Supplementary Figure S4. The position (viral protein (VP) and residue number) and location of the identified cleavage sites are listed. The mutants are referred to as pE11.VPn.XmZ, where amino acid X at position m in viral protein n is replaced by amino acid Z. E: echovirus, EV-B: *Enterovirus betacoxsackie*, CV: coxsackievirus.

| Selected residue | Location on viral capsid | Name of mutant | Enterovirus types with the substituted amino acid |
| --- | --- | --- | --- |
| VP1.K155 | EF loop | pE11.VP1.K155N | E3, E7, E12, EV-B101 |
| VP1.K274 | C terminus | pE11.VP1.K274E | E3, E7, CVB4 |
| VP2.Y97 | CD loop | pE11.VP2.Y97Q | CVA9, CVB1, CVB3, CVB4, CVB5 |
| VP3.R69 | BC loop | pE11.VP3.R69Q | CVB3, CVB5, EV-B97 |

### Assessing the inactivation of E11 mutants in lakewater

To assess the contribution of each proposed proteolytic cleavage site to E11 inactivation in lakewater, we introduced targeted point mutations at these sites using reverse genetics (**Table 1**). Each amino acid substitution (VP1.K155N, VP1.K274E, VP2.Y97Q, and VP3.R69Q) was chosen based on naturally occurring amino acids present in other enterovirus types. We incubated the plasmid-produced E11 wild-type (pE11) and the mutant viruses in lakewater and monitored their infectivity over time (**Figure 3**). The plasmid-produced virus rather than the E11 wild-type was used here to ensure comparability with the mutant virus propagation method, and this virus exhibited inactivation kinetics indistinguishable from those of the wild-type (analysis of covariance: p = 0.28, **Supplementary Figure S6**).

**Figure 3.**
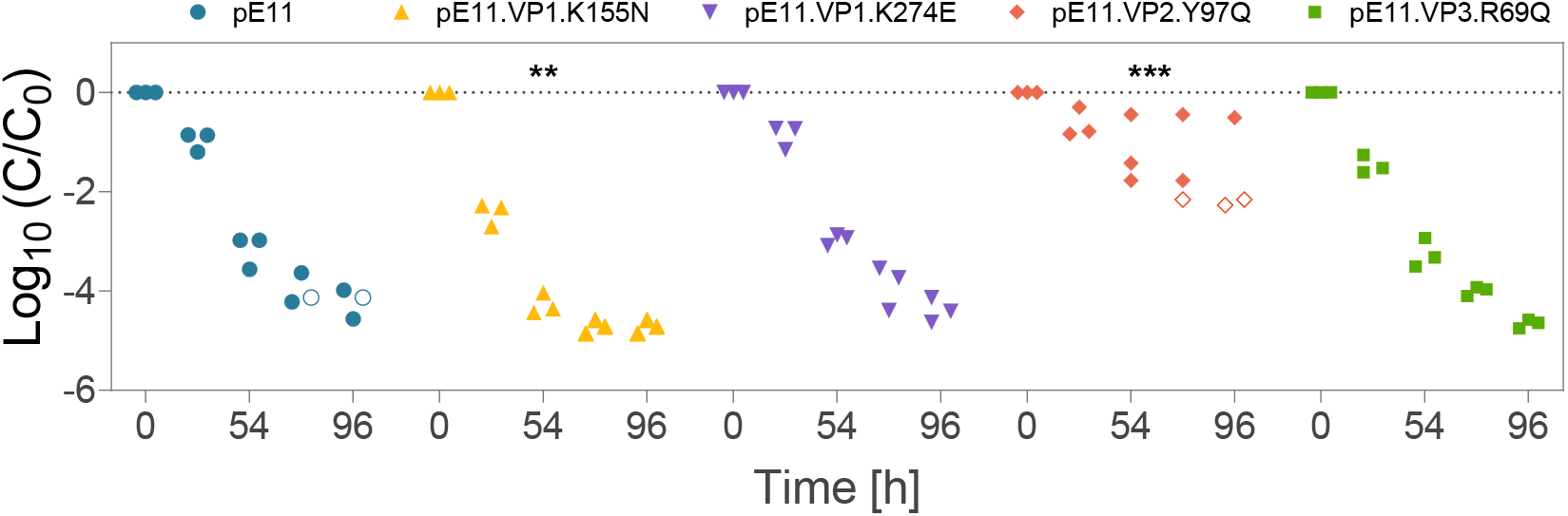
The inactivation of the plasmid-produced E11 wild-type (pE11) and four mutant viruses in surface water from Lake Geneva. Lakewater was collected in September 2025. All viruses were added to an initial concentration of 10^6^ MPNCU/mL except for mutant pE11.VP2.Y97Q which was added to 10^4^ MPNCU/mL. Individual data points of triplicate experiments are presented. Values below the detection limit are indicated by open symbols and set to LOQ/√2 where LOQ = 90.4 MPNCU/mL is the limit of detection. Values below the LOQ were not used for calculation of inactivation rate constants (*k*, **Supplementary Table S1**). Note that the different measurable range in inactivation results from differences in the starting titers. Significant differences in mutant decay with respect to wild-type decay were determined by analysis of covariance using Tuckey’s honest significant difference method to correct for multiple comparisons (**: p < 0.01; ***: p < 0.001).

Among the mutants, we observed that pE11.VP2.Y97Q exhibited markedly increased stability, with an 11-fold lower inactivation rate constant than the wild-type (pE11.VP2.Y97Q: *k* = 0.01 ± 0.01 h^-1^; pE11: *k* = 0.11 ± 0.01 h^-1^, **Supplementary Table S1**), indicating that cleavage C-terminal to residue VP2.Y97 is a key driver of E11 inactivation. In contrast, mutants pE11.VP1.K274E and pE11.VP3.R69Q displayed inactivation rates indistinguishable from pE11 (all *k* = 0.11 ± 0.01 h^-1^) whereas pE11.VP1.K155N decayed 1.6-fold more rapidly (*k* = 0.18 ± 0.02 h^-1^). These results indicate that residues VP1.K155, VP1.K274, and VP3.R69 do not substantially contribute to the proteolytic inactivation of E11. Additionally, we found that all mutants were more stable in sterile rather than in microbially active lakewater (**Supplementary Figure S7**), indicating that mutagenesis did not compromise viral viability and that microbial activity promotes inactivation of the mutant viruses.

To exclude the potential effect of initial virus concentration on inactivation kinetics, we performed a confirmatory experiment in which we added all viruses to the same initial titer (10^4^ most probable number of cytopathic units per milliliter (MPNCU/mL)). We again observed greater stability of mutant pE11.VP2.Y97Q relative to pE11 (**Supplementary Figure S8**), confirming the essential role of residue VP2.Y97 in governing the proteolytic inactivation of E11 in lakewater.

To further evaluate proteolytic cleavage adjacent to the proposed sites, we assessed E11 stability in the presence of trypsin and chymotrypsin, which cleave C-terminal to lysine and arginine residues, and to aromatic residues including tyrosine, respectively^28^ (**Supplementary Figure S9A**). We found that E11 remained stable in the presence of trypsin (**Supplementary Figure S9B**), consistent with the lack of contribution of residues VP1.K155, VP1.K274, and VP3.R69 to viral inactivation. Interestingly, E11 also remained stable in the presence of chymotrypsin (**Supplementary Figure S9B**). These results indicate that the protease(s) responsible for cleavage proximal to residue VP2.Y97 in lakewater do not share the substrate specificity of chymotrypsin and likely require additional sequence or structural determinants beyond the P1 residue.

### Capsid amino acid composition governs the stability of enteroviruses in lakewater

Lastly, we compared E11 decay in lakewater with that of echovirus 3 (E3), echovirus 7 (E7), and CVB5. We selected these enteroviruses based on amino acid variation at the proposed E11 cleavage sites (**Table 2 & Supplementary Figures S10-12**). Sequence analysis showed that CVB5 lacks all four proposed E11 target residues, whereas E3 and E7 retain only VP3.R69 and VP1.K274, respectively. Notably, residue VP2.Y97 was absent from all three viruses. Across viral proteins, sequence identity relative to E11 ranged from 69% to 84% identity (**Supplementary Figures S10-12**).

**Table 2.**
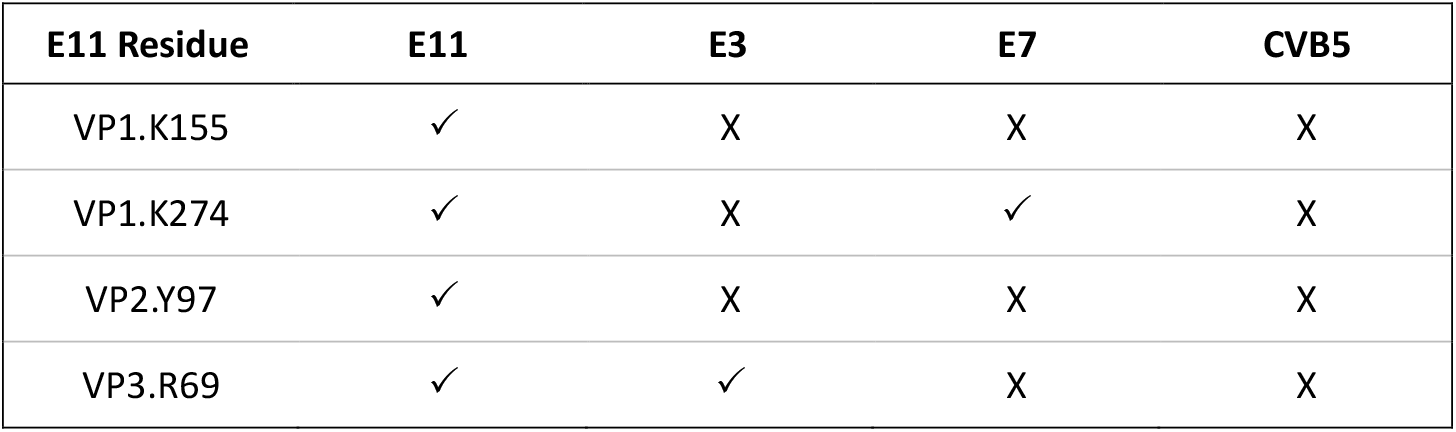
Difference in amino acid composition between the *Enterovirus betacoxsackie* types studied here. Presence (✓) or absence (X) of the four proposed E11 proteolytic target sites in the viral proteins of E11, E3, E7, and CVB5.

We observed marked differences in stability among the enterovirus types in lakewater (**Figure 4**). E11 was rapidly inactivated, whereas E3 retained infectivity for approximately 30 hours before a pronounced decline, and both E7 and CVB5 remained stable for at least 54 hours. The increased stability of E3, E7, and CVB5 relative to E11 was consistent with the absence of VP2.Y97 in these enteroviruses. In addition, the stability of E7 supported the lack of a detectable contribution of VP1.K274 to viral decay. The delayed loss of infectivity observed for E3 suggests a temporally distinct inactivation process under the same conditions, perhaps due to microbial growth in lakewater leading to the delayed secretion of E3-specific proteases. Alternatively, E3 may undergo multi-hit decay^33^, requiring more proteolytic cleavage events to initiate inactivation compared with E11.

**Figure 4.**
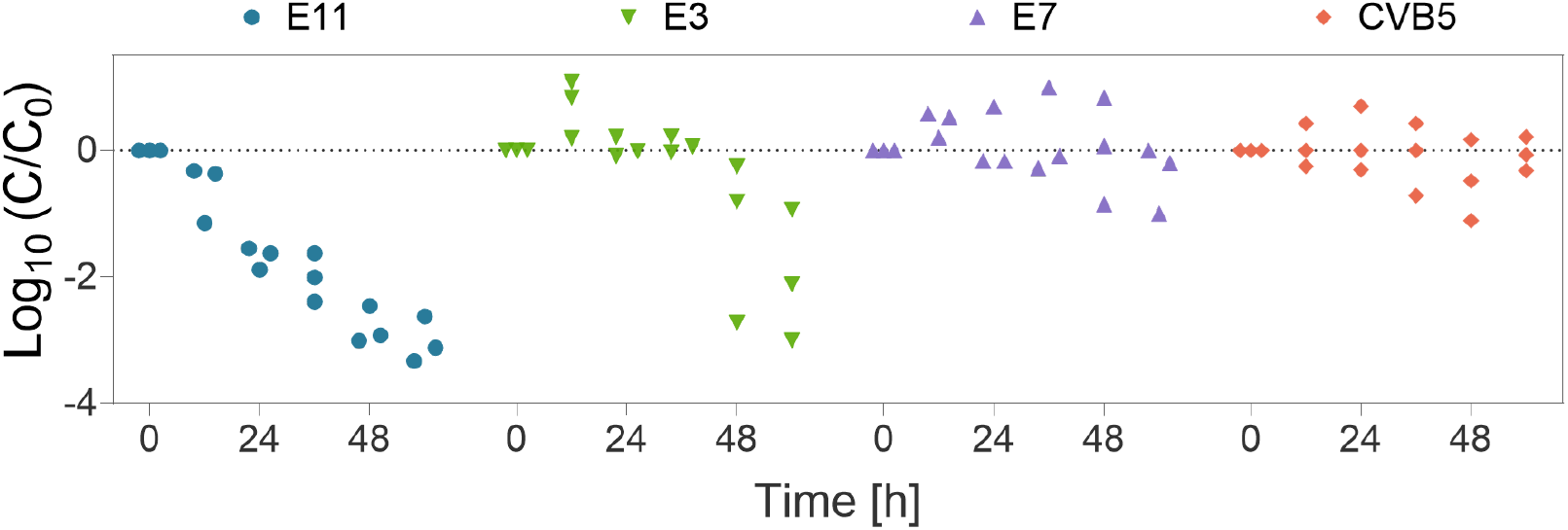
The inactivation of four *Enterovirus betacoxsackie* types, E11, E3, E7, and CVB5, in surface water from Lake Geneva collected in April 2025. Individual data points of triplicate experiments are presented.

Overall, these findings suggest that the amino acid composition of viral proteins is essential for governing the fate of enteroviruses in lakewater.

## Discussion

In accordance with earlier reports^13,18^, we observed rapid inactivation of E11 in microbially active lakewater (**Figure 1A & Supplementary Figure S1**), although variability in the antiviral activity of lakewater was noted. We found that metalloproteases contributed substantially to this antiviral activity (**Supplementary Figure S2**), supporting a protease-mediated mechanism. Because other protease classes, including serine proteases, have been previously implicated in enterovirus inactivation^14,18^, the total contribution of proteases to E11 decay in lakewater is likely underestimated here.

The Lake Geneva protease pool exhibited variable antiviral activity and heterogeneity in substrate specificity, yet we observed that the dominant features of the proteolytic fingerprints remained largely conserved (**Supplementary Figure S3**). These conserved features were used to identify four candidate proteolytic cleavage sites on the surface of E11. Using site-directed mutagenesis, we showed that cleavage proximal to residue VP2.Y97 contributes substantially to E11 inactivation in lakewater (**Figure 3**). In contrast, we found that substitution of the three remaining target residues had no detectable effect on virus stability, indicating that cleavage at these sites does not drive E11 decay. Nevertheless, we cannot fully exclude proteolysis at these positions, as cleavage may not induce sufficient structural disruption of the capsid to result in loss of infectivity. The observed delay in E3 decay (**Figure 4**) may result from cleavage at residue VP3.R69, which is shared with E11 but not with any of the other enteroviruses tested. Variations in capsid structure and sequence composition between these two viral pathogens may therefore account for the observed difference in stability.

The minimal contribution of residues VP1.K155, VP1.K274, and VP3.R69 to E11 inactivation suggests that the prevailing trypsin-like specificity of the protease pool – characterized by cleavage C-terminal to lysine or arginine^28^ - is unlikely to account for E11 decay. We supported this conclusion by showing that E11 remained stable in the presence of trypsin (**Supplementary Figure S9**). Furthermore, we found that, with the exception of residue VP1.K155, none of the proposed cleavage sites interact with either the complement decay-accelerating factor (CD55) attachment receptor or the human neonatal Fc (FcRn) uncoating receptor used by E11 for host cell entry^34^. We therefore hypothesize that cleavage at VP2.Y97 is unlikely to impair receptor binding or cell entry. Instead, we propose that cleavage at VP2.Y97 may induce local unraveling and instability of VP2, thereby exposing additional sites to subsequent proteolytic attack and ultimately leading to capsid damage^31^.

Notably, we found that residue VP2.Y97 is absent from the viral protein sequences of E3, E7 and CVB5 (**Table 2**), where it is replaced by glutamic acid, leucine, and glutamine, respectively (**Supplementary Figure S11**). Accordingly, these enterovirus types exhibited high stability in lakewater (**Figure 4**), suggesting that the role of VP2.Y97 may extend to other enterovirus types. This insight is particularly relevant for understanding the fate of viruses that are difficult to culture, such as EV-A71 and EV-D68, or to test due to containment policies, such as poliovirus. EV-A71 and EV-D68 - both associated with severe neurological infections^35,36^ - also lack VP2.Y97 and may therefore exhibit enhanced environmental stability relative to E11.

Although we identify residue VP2.Y97 as a primary determinant of enterovirus susceptibility to environmental proteolysis, it is unlikely to be the sole contributor. We anticipate that sequence and structural variation across enterovirus capsids may generate alternative protease-sensitive sites, modulating environmental stability in a virus-specific manner. In this study, we identified candidate target sites based exclusively on the dominant substrate specificities of the lakewater protease pool. Because the proteolytic fingerprints capture only significantly enriched or depleted residues flanking detected cleavage sites, rare or kinetically slow proteolytic events are not represented in the analysis. Nevertheless, we acknowledge that such low-frequency cleavage events may still contribute to viral inactivation if they occur at structurally critical regions of the capsid. Consistent with this possibility, we observed that increased heterogeneity in substrate specificity - reflected by a greater number of detected cleavage sites - was associated with more rapid E11 decay (**Figure 1C**). We therefore identify minor proteolytic substrate specificities as an important avenue for future investigation.

E11 is a ubiquitous contaminant of surface waters and is highly relevant in clinical settings, where infections may prove fatal to newborns. Efficient transmission of this pathogen through contaminated waters to mothers thus poses a potential risk during pregnancy. However, the presence of residue VP2.Y97 in the E11 isolate used in this study and in variants isolated from neonatal clinical specimens^4^ suggests that this virus is intrinsically labile in the environment. Such instability is noteworthy, as reduced survival in natural waters may constrain transmission despite the high prevalence of E11 in human populations.

Collectively, these findings highlight the central role of capsid amino acid composition in shaping enterovirus stability in natural waters. By coupling the substrate specificity of an environmental protease pool with viral protein amino acid composition and structural features, we demonstrate a framework that may enable prediction of enterovirus environmental fate and explain why closely related enteroviruses exhibit markedly different stability outside the host. We anticipate that this approach can be used to anticipate environmental stability across diverse enterovirus types and will be particularly valuable for enteroviruses that are difficult to culture or experimentally assess.

## Methods

### Virus propagation, enumeration, and sequencing

Echovirus 11 (E11, propagated from Gregory strain ATCC VR737), echovirus 3 (E3, propagated from an environmental isolate provided by the Finnish National Institute for Health and Welfare), and echovirus 7 (E7, propagated from an environmental isolate provided by the Finnish National Institute for Health and Welfare) stocks were prepared by infecting sub-confluent layers of Rhabdomyosarcoma cells (RD, ATCC CCL-136) in T-75 culture flasks. The cells were grown and maintained at 37°C and 5% CO_2_ with Dulbecco’s Modified Eagle Medium (DMEM, 41966, Gibco) supplemented with 1% penicillin-streptomycin (15140, Gibco) and 10% (growth) or 2% (maintenance) heat-inactivated fetal bovine serum (FBS, A5256701, Gibco). Similarly, coxsackievirus B5 (CVB5, propagated from an environmental isolate, GenBank accession number MG845891) was propagated in Buffalo green monkey kidney cells (BGMK, kindly provided by Spiez Laboratory, Switzerland) with identically penicillin-streptomycin and FBS-supplemented minimum essential medium (MEM, A41922). The viruses were released from the cells three days post infection by freeze-thawing the culture flasks three times. Cell debris was removed by centrifugation at 1’100x *g* for 5 min and buffer-exchanged into phosphate-buffered saline (PBS, 18912014, Gibco) using Amicon Ultra-15 centrifugal filters (UFC910024, Merck Millipore).

Infectious E11, E3, E7, and CVB5 virions were enumerated using a most probable number (MPN) infectivity assay as described previously^37^. Briefly, RD or BGMK cells were grown to 95% confluence in 96-well plates (Greiner CELLSTART, Sigma Aldrich). Then, 20 µL of each sample (in five replicates) were 10-fold serially diluted in maintenance medium and added to the cells. The plates were incubated at 37°C and 5% CO_2_ for 5 days after which time each well was visually examined for cytopathic effect (CPE) using an inverted microscope. The infectious virus titer was determined as the MPN of cytopathic units per milliliter (MPNCU/mL) by converting the number of wells positive for CPE using the R package *MPN*^38^. The assay limit of quantification (LOQ) was considered as one positive well (of five replicates) at the highest dilution (10-fold diluted) and corresponded to 90.4 MPNCU/mL. Values below the LOQ were set to LOQ/√2 and indicated by open symbols in all plots. The coefficient of variation of enumeration was determined to be 40-50 % based on replicate titration of E11 (n = 12) and CVB5 (n = 9). The titers of the obtained virus stocks were 10^8^ or 10^11^ MPNCU/mL for E11, 10^9^ MPNCU/mL for CVB5 and E3, and 10^10^ MPNCU/mL for E7.

E11, E3, E7, and CVB5 structural proteins were sequenced using the primer pairs and cycling parameters listed in **Supplementary Tables S3 and S4**. Details regarding virus sequencing can be found in the Supplementary Methods. Viral protein 2 (VP2) of E7 could only be partially sequenced. The full genome of E11 and full or partial viral protein sequences of E3, E7, and CVB5 are listed in the Supplementary Methods. Using the determined sequence of E11, the viral capsid was predicted and modeled using AlphaFold2-Multimer1^39,40^ in ColabFold3^41^ and ChimeraX^32^. Details regarding the structural representation of E11 can be found in the Supplementary Methods.

### Design and production of E11 mutants by reverse genetics

Echovirus 11 mutants were designed based on naturally occurring amino acid substitutions found in closely-related enterovirus types at the proposed proteolytic cleavage sites. Briefly, GenBank accession numbers of enterovirus types were obtained from the Virus Metadata Resource (version VMR_MSL38_v2) from the International Committee on Taxonomy of Viruses^42^. The protein sequences of the identified isolates were compiled and aligned in Geneious Prime 2025.1.3 (https://www.geneious.com) by Multiple Alignment using Fast Fourier Transform (MAFFT)^43^. A neighbor-joining phylogenetic tree was subsequently computed and the enterovirus types presenting strong homology to E11 were used to list possible amino acid substitutions for each proposed cleavage site (**Supplementary Table S5**). The final amino acid substitutions (**Table 1**) were chosen based on the following criteria: (i) the residue is not favored for proteolytic cleavage as determined by MSP-MS and (ii) the surface charge and hydrophobicity of the residue are similar to those of the residue to be substituted.

Infectious cDNA clones for wild-type E11 and E11 mutants were constructed by Genscript Biotech. Sequences of T7 promotor followed by full-length viral genome, polyA, and MluI restriction sites were cloned into a pUC57 vector. The full length wild-type and mutant RNA genomes were produced by *in vitro* transcription of the MluI-linearized plasmids using the RiboMAX T7 Express Large Scale RNA Production System (P1320, Promega). Subsequently, 2-3 µg transcribed RNA genome was transfected into 70-90% sub-confluent BGMK cells using Lipofectamine MessengerMAX (LMRNA001, Thermo Fisher) as a transfection reagent. Upon onset of full CPE (3-4 days), virus progeny was collected and purified as described above and the amino acid substitutions in the mutant E11 viruses were confirmed by Sanger sequencing. Details regarding the production of E11 wild-type and mutants by reverse genetics can be found in the Supplementary Methods. The full genome of the infectious E11 wild-type and mutated cDNA clones were deposited under GenBank accession numbers PX993811, PX993812, PX993813, PX993814, and PX993815. The E11 wild-type and mutant viruses are referred to as pE11 and pE11.VPn.XmZ respectively, where amino acid X at position m in the viral protein n is replaced by amino acid Z. The titers of the obtained virus progeny stocks were 10^8^ MPNCU/mL for pE11 and pE11.VP1.K155N, 10^9^ MPNCU/mL for pE11.VP1.K274E and pE11.VP3.R69Q, and 10^5^ MPNCU/mL for pE11.VP2.Y97Q.

### Lakewater sampling and processing

Surface lakewater was collected from Lake Geneva on April 9^th^ 2024, July 23^rd^ 2024, September 10^th^ 2024, October 1^st^ 2024, February 3^rd^ 2025, April 14^th^ 2025, July 21^st^ 2025, and September 15^th^ 2025 from the shore in Saint-Sulpice, Switzerland or from the floating research platform LéXPLORE located 570 m off the shore of Pully, Switzerland (October 2024 sample, https://lexplore.info/). Immediately after collection, the lakewater samples were brought to the laboratory and vacuum-filtered through a 0.8 μm MCE membrane filter (AAWP04700, MF-Millipore) to remove eukaryotes while retaining bacteria in the lakewater^44^. The resulting filtrate was termed microbially active lakewater. Sterile lakewater was obtained by autoclaving the filtered lakewater at 121°C for 15 minutes. Both microbially active and sterile lakewater samples were stored at 4°C overnight prior to virus inactivation experiments and determination of the proteolytic fingerprint.

### Virus inactivation experiments in lakewater

Inactivation experiments with E11 were conducted in all lakewater samples and those of CVB5, E3, and E7 in lakewater collected in April 2025. The inactivation of E11 mutants in lakewater was studied in August and September 2025. Briefly, 1-10 mL of microbially active and sterile lakewater samples were spiked with virus to an initial titer of approximately 10^7^ MPNCU/mL and infectivity was monitored over time. For E11 mutant inactivation experiments, an initial titer of 10^6^ MPNCU/mL was used for wild-type pE11 and all E11 mutants except for pE11.VP2.Y97Q, for which an initial titer of 10^4^ MPNCU/mL was used due to the low titer of the progeny virus. Aliquots of 150 μL were taken at timepoints across 54 hours and stored at -20°C until virus enumeration by MPN infectivity assay. The samples were maintained shaking gently at room temperature (KS 501 digital shaker, IKA) throughout the experiment. Each experiment was conducted in triplicate. In experiments where several viruses were studied, the viruses were added individually into separate lakewater samples. Virus inactivation rate constants (*k*, h^-1^) were determined by least square fit of inactivation data pooled across triplicate experiments to a first-order decay model (Equation 1):

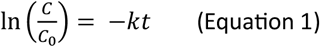

where *C* is the infectious virus titer at time *t* (h) and *C*_*0*_ is in initial infectious virus titer. Data points below the LOQ were not considered for the determination of rate constants. Values of *k* are reported with 95% confidence intervals associated with the regression slope.

### Determining the proteolytic fingerprint of lakewater by Multiplex Substrate Profiling by Mass Spectrometry (MSP-MS)

The proteolytic fingerprints of all Lake Geneva samples were determined using the MSP-MS protocol devised by O’Donoghue *et al*.^20,45^ with an adapted enzyme quenching process^46^. Briefly, the MSP-MS peptide library was spiked into 500 μL microbially active or sterile lakewater (in triplicate) to a final peptide concentration of 0.5 μM per peptide. After 6 hours, aliquots of 55 μL were heat treated at 80°C (water bath) for 10 minutes to inactivate the proteases. The inactivated samples were centrifuged for 1 minute at maximum speed (25’000x *g*) and the supernatant (50 μL) was collected. The samples were stored at -20°C until further analysis by liquid chromatography coupled to high-resolution mass spectrometry (LC-HRMS/MS)^27^. The eight residues (P4-P4’) surrounding the identified cleavage sites (P1↓P1’) were used to generate the proteolytic fingerprint of the sample in the form of an iceLogo plot^47^. The iceLogo software calculates the percent difference between the frequencies of an amino acid in the detected cleavage products and the MSP-MS peptide library at each position. Only significantly under- or over-represented residues (t-test, p ≤ 0.05) are included in the iceLogo plots. Details regarding MSP-MS peptide library composition and preparation, LC-HRMS/MS running parameters, and data analysis have been previously described^27^.

### Metalloprotease inhibitor experiment

The effect of the metalloprotease inhibitor GM6001 (CC1010, Sigma Aldrich) on E11 inactivation was studied in July 2024 and July 2025. GM6001 was added into microbially active lakewater at a concentration of 5 μM and incubated at room temperature for 30 min prior to performing virus inactivation experiments.

### Statistical analysis

Statistical comparison of data from two groups was performed using a t-test while comparison of multiple groups was performed by one-way analysis of variance (ANOVA) with Dunnett’s test for multiple comparisons assuming normally distributed data. Analysis of covariance (ANCOVA) was used for comparison of decay kinetics between multiple groups using Tuckey’s honest significant difference method to correct for multiple comparisons. For assessment of the correlation between two variables, Pearson’s correlation coefficient (r) was calculated assuming a normal distribution. Analysis of covariance was performed using the *emmeans* R package^48^. All other analyses were conducted using GraphPad Prism v10.6.1 (GraphPad Software, USA).

## Acknowledgements

This work was funded by the Swiss National Science Foundation (grant no. 310030 215226). ST is supported by the JSPS through for Young Scientists (grant number 24K17379), Kurita Water and Environmental Foundation (grant number 23H036, 24K023, and 25K023). MZ acknowledges funding by the Austrian Science Fund (FWF), [Cluster of Excellence CoE7, Grant DOI: 10.55776/COE7].

## Data Availability

All data are available on Zenodo under link https://doi.org/10.5281/zenodo.18545921. Full genomes of virus and infectious clones are available at NCBI under accession numbers PX993811, PX993812, PX993813, PX993814, and PX993815.

## References

1. Harvala, H. et al. Enterovirus circulation in the WHO European region, 2015–2022: a comparison of data from WHO’s three core poliovirus surveillance systems and the European Non-Polio Enterovirus Network (ENPEN). The Lancet Regional Health - Europe 53, 101292 (2025).

2. de Schrijver, S. et al. Epidemiological and Clinical Insights into Enterovirus Circulation in Europe, 2018–2023: A Multicenter Retrospective Surveillance Study. J Infect Dis 232, e104–e115 (2025).

3. Vlok, M. & Majer, A. Global Prevalence of Non-Polio Enteroviruses Pre- and Post COVID-19 Pandemic. Microorganisms 13, 1801 (2025).

4. Grapin, M. et al. Severe and fatal neonatal infections linked to a new variant of echovirus 11, France, July 2022 to April 2023. Eurosurveillance 28, 2300253 (2023).

5. European Centre for Disease Prevention and Control. Epidemiological update: Echovirus 11 infections in neonates. https://www.ecdc.europa.eu/en/news-events/epidemiological-update-echovirus-11-infections-neonates (2023).

6. Ikuse, T. et al. Neonatal acute liver failure cases with echovirus 11 infections, Japan, August to November 2024. Euro Surveill 30, 2400822 (2025).

7. Fang, C., Zhang, X., Huang, X., Xu, F. & Zhao, D. Epidemiology and control measures of an outbreak of neonatal echovirus 11 infections in Guangdong, China: A retrospective analysis. Biosafety and Health 5, 227–232 (2023).

8. Fong, T.-T. & Lipp, E. K. Enteric Viruses of Humans and Animals in Aquatic Environments: Health Risks, Detection, and Potential Water Quality Assessment Tools. Microbiol Mol Biol Rev 69, 357–371 (2005).

9. European Centre for Disease Prevention and Control. Disease information on human non-polio enterovirus infections. https://www.ecdc.europa.eu/en/enteroviruses/facts (2025).

10. Fujioka, R. S., Loh, P. C. & Lau, L. S. Survival of human enteroviruses in the Hawaiian Ocean environment: Evidence for virus-inactivating microorganisms. Applied and Environmental Microbiology 39, 1105–1110 (1980).

11. Ward, R. L., Knowlton, D. R., Winston, P. E. & Gamble, J. N. Mechanism of Inactivation of Enteric Viruses in Fresh Water. Applied and Environmental Microbiology 52, 450–459 (1986).

12. Gordon, C. & Toze, S. Influence of groundwater characteristics on the survival of enteric viruses. Journal of Applied Microbiology 95, 536–544 (2003).

13. Romanenko, A., Peter, H., Meibom, J., Borchardt, M. A. & Kohn, T. Diversity of lake bacteria promotes human echovirus inactivation. Applied and Environmental Microbiology 91, e02366–24 (2025).

14. Cliver, D. O. & Herrmann, J. E. Proteolytic and Microbial Inactivation of Enteroviruses. Water Research 6, 797–805 (1972).

15. Qin, Y. et al. Purification and Characterization of a Secretory Alkaline Metalloprotease with Highly Potent Antiviral Activity from Serratia marcescens Strain S3. Journal of Agricultural and Food Chemistry 67, 3168–3178 (2019).

16. Corre, M.-H. et al. The early communication stages between serine proteases and enterovirus capsids in the race for viral disintegration. Commun Biol 7, 969 (2024).

17. Yamamoto, S. et al. Bacillaceae serine proteases and Streptomyces epsilon-poly-l-lysine synergistically inactivate Caliciviridae by inhibiting RNA genome release. Scientific Reports 14, (2024).

18. Corre, M. H., Bachmann, V. & Kohn, T. Bacterial matrix metalloproteases and serine proteases contribute to the extra-host inactivation of enteroviruses in lake water. ISME Journal 16, 1970–1979 (2022).

19. Myouga, H., Yoshimizu, M., Tajima, K. & Ezura, Y. Purification of an Antiviral Substance Produced by Alteromonas sp. and Its Virucidal Activity against Fish Viruses. Fish Pathology 30, 15–22 (1995).

20. O’Donoghue, A. J. et al. Global identification of peptidase specificity by multiplex substrate profiling. Nature Methods 9, 1095–1100 (2012).

21. Wichmann, N., Gruseck, R. & Zumstein, M. Hydrolysis of Antimicrobial Peptides by Extracellular Peptidases in Wastewater. Environmental Science and Technology 58, 717–726 (2024).

22. Wichmann, N., Meibom, J., Kohn, T. & Zumstein, M. Conserved specificity of extracellular wastewater peptidases revealed by multiplex substrate profiling by mass spectrometry. Environ Chem Lett 23, 953–959 (2025).

23. Kimura, T., Yoshimizu, M. U., Ezura, Y. & Kamel, Y. An Antiviral Agent (46NW-04A) Produced by Pseudomonas sp. and Its Activity against Fish Viruses. Journal of Aquatic Animal Health 2, 12–20 (1990).

24. Carpine, R. & Sieber, S. Antibacterial and antiviral metabolites from cyanobacteria: Their application and their impact on human health. Current Research in Biotechnology 3, 65–81 (2021).

25. Padhi, C. et al. Metagenomic study of lake microbial mats reveals protease-inhibiting antiviral peptides from a core microbiome member. Proc. Natl. Acad. Sci. U.S.A. 121, e2409026121 (2024).

26. Kamei, Y., Yoshimizu, M., Ezura, Y. & Kimura, T. Screening of bacteria with antiviral activity from fresh water salmonid hatcheries. Microbiol Immunol 32, 67–73 (1988).

27. Meibom, J., Wichmann, N., Astorch-Cardona, A., Zumstein, M. & Kohn, T. Proteolytic Activity and Substrate Specificity of Lake Geneva. Environ. Sci. Technol. 59, 27811–27823 (2025).

28. Harris, J. L. et al. Rapid and general profiling of protease specificity by using combinatorial fluorogenic substrate libraries. Proceedings of the National Academy of Sciences U.S.A 97, 7754–7759 (2000).

29. Niu, S. et al. Molecular and structural basis of Echovirus 11 infection by using the dual-receptor system of CD55 and FcRn. Chinese Science Bulletin 65, 67–79 (2020).

30. Fontana, A. et al. Correlation between sites of limited proteolysis and segmental mobility in thermolysin. Biochemistry 25, 1847–1851 (1986).

31. Fontana, A. et al. Probing protein structure by limited proteolysis. Acta Biochim Pol 51, 299–321 (2004).

32. Pettersen, E. F. et al. UCSF ChimeraX: Structure visualization for researchers, educators, and developers. Protein Sci 30, 70–82 (2021).

33. Gyürék, L. L. & Finch, G. R. Modeling Water Treatment Chemical Disinfection Kinetics. Journal of Environmental Engineering 124, 783–793 (1998).

34. Zhao, X. et al. Human Neonatal Fc Receptor Is the Cellular Uncoating Receptor for Enterovirus B. Cell 177, 1553–1565.e16 (2019).

35. Ooi, M. H., Wong, S. C., Lewthwaite, P., Cardosa, M. J. & Solomon, T. Clinical features, diagnosis, and management of enterovirus 71. The Lancet Neurology 9, 1097–1105 (2010).

36. Messacar, K. et al. A cluster of acute flaccid paralysis and cranial nerve dysfunction temporally associated with an outbreak of enterovirus D68 in children in Colorado, USA. The Lancet 385, 1662–1671 (2015).

37. Carratalà, A. et al. Experimental adaptation of human echovirus 11 to ultraviolet radiation leads to resistance to disinfection and ribavirin. Virus Evolution 3, vex035 (2017).

38. Ferguson, M. & Ihrie, J. MPN: Most Probable Number and Other Microbial Enumeration Techniques. (2024).

39. Jumper, J. et al. Highly accurate protein structure prediction with AlphaFold. Nature 596, 583–583 (2021).

40. Evans, R. et al. Protein complex prediction with AlphaFold-Multimer. 2021.10.04.463034 Preprint at 10.1101/2021.10.04.463034 (2022).

41. Mirdita, M. et al. ColabFold: making protein folding accessible to all. Nature Methods 19, 679–682 (2022).

42. The International Committee on Taxonomy of Viruses (ICTV): Virus Metadata Resource (VMR). https://ictv.global/vmr.

43. Katoh, K., Misawa, K., Kuma, K. & Miyata, T. MAFFT: a novel method for rapid multiple sequence alignment based on fast Fourier transform. Nucleic Acids Res 30, 3059–3066 (2002).

44. Olive, M., Gan, C., Carratalà, A. & Kohn, T. Control of waterborne human viruses by indigenous bacteria and protists is influenced by temperature, virus type, and microbial species. Applied and Environmental Microbiology 86, e01992–19 (2020).

45. Rohweder, P. J., Jiang, Z., Hurysz, B. M., O’Donoghue, A. J. & Craik, C. S. Multiplex substrate profiling by mass spectrometry for proteases. Methods in Enzymology 682, 375–411 (2023).

46. Wichmann, N., Meibom, J., Kohn, T. & Zumstein, M. Conserved specificity of extracellular wastewater peptidases revealed by multiplex substrate profiling by mass spectrometry. Environ Chem Lett 10.1007/s10311-025-01834-7 (2025) doi:10.1007/s10311-025-01834-7.

47. Colaert, N., Helsens, K., Martens, L., Vandekerckhove, J. & Gevaert, K. Improved visualization of protein consensus sequences by iceLogo. Nature Methods 6, 786–787 (2009).

48. Lenth, R. V. et al. emmeans: Estimated Marginal Means, aka Least-Squares Means. (2025).

