## Supplementary Methods and Information for "A Capsid Tyrosine Residue Governs the Proteolytic Inactivation of Echovirus 11 in Lakewater"

Number of pages: 22

Number of tables: 5

Number of figures: 12

#### Supplementary Methods

*Virus sequencing*

To obtain the full genome sequence of E11, reverse transcription (RT)-PCR followed by Sanger sequencing was perfromed. Similarly, the viral proteins (VP1-3) of E3, E7, and CVB5 were sequenced. Briefly, viral RNA was extracted from virus stocks using the QIAamp viral RNA Mini Kit (52906, Qiagen) according to the manufacturer's instructions. Subsequently, amplicons were produced using the SuperScript IV UniPrime One-Step RT-PCR System (12596025, Thermo Fisher) with the primer pairs listed in **Supplementary Table S3**. The RT-PCR reaction volumes were 50 µL and each consisted of 25 µL 2x UniPrime RT-PCR Master Mix, 2.5 µL each of forward and reverse primers (10 µM), 2 µL 25x SuperScript IV RT Mix, 5 µL RNA template, and 13 µL nuclease-free water. RT-PCR cycling parameters for each primer pair are listed in **Supplementary Table S4**.

*Production of E11 mutants by reverse genetics*

Infectious cDNA clones for wild-type E11 and E11 mutants were constructed by Genscript Biotech. Sequences of T7 promotor followed by full-length viral genome, polyA, and MluI restriction sites were cloned into a pUC57 vector. The plasmids were propagated in competent *E. coli* cells (DH5α, C2987H, NEB) and purified using NucleoBond Xtra Midi columns (740410.1, Takara Bio) according to the manufacturers' instructions. Following linearization of the plasmids using MluI restriction enzyme (R3198S, NEB), full-length E11 RNA genomes were produced by *in vitro* transcription using the RiboMAX T7 Express Large Scale RNA Production System (P1320, Promega) and transfected into 70-90% sub-confluent BGMK cells in T-25 culture flasks. Specifically, 5.5 µL transfection reagent (Lipofectamine MessengerMAX, LMRNA001, Thermo Fisher) and 2-3 µg transcribed RNA genome were incubated separately in 250 µL Opti-MEM medium (31985062, Gibco) for 10 min at room temperature after which time the two solutions were mixed and further incubated for 5 min. Then, the RNA-lipofectamine mixture was used to transfect the BGMK cells which were incubated in MEM maintenance medium at 37°C and 5% $CO_2$ for 3-4 days until full CPE was observed. Finally, virus progeny were collected and purified (see main text) and the amino acid substitutions in the mutant E11 viruses were confirmed by Sanger sequencing.

*Structural representation of E11*

Using AlphaFold2-Multimer[1,2] (AF2-M) v3 in ColabFold[3] v1.5.5, the structure of the E11 protomer (VP1-3). Specifically, multiple sequence alignments (MSA) were generated with MMseqs2[4] against the UniRef100[5] and environmental databases using the unpaired_paired mode. Structures in the PDB100 database[6] was used as templates for the prediction. The folding of the protomer was performed using

3 recycles. The quality of the predicted protomer model was assessed using the predicted local Distance Difference Test (pLDDT), predicted Template Modeling (pTM), interface predictive Template Model (ipTM), and Predicted Alignment Error (PAE) scores (**Supplementary Figure S5**). The overall model confidence was calculated using Equation S1:

$$\text{Model confidence} = 0.8 \times \text{ipTM} + 0.2 \times \text{pTM} \qquad \text{(Equation S1)}$$

The overall model confidence was calculated to be 0.939.

Using the first ranked predicted protomer model, the full virus capsid of E11 was constructed in UCSF ChimeraX[7] v1.10.1 by docking the modeled protomer onto that of an experimentally resolved E11 structure (PDB 6LA3[8]) and applying icosahedral symmetry. Each amino acid substitution relative to the sequence of PDB 6LA3 was individually evaluated for potential clashes within the structure. The predicted protomer model and the constructed virus capsid of E11 were used to produce visual representations in ChimeraX.

*E11 plasmid and mutant sequences*

>pE11G

TCGCGCGTTTCGGTGATGACGGTGAAAACCTCTGACACATGCAGCTCCCGGAGACGGTCACAGCTTGTCTGTAAGCGGATGC
CGGGAGCAGACAAGCCCGTCAGGGCGCGTCAGCGGGTGTTGGCGGGTGTCGGGGCTGGCTTAACTATGCGGCATCAGAGC
AGATTGTACTGAGAGTGCACCATATGCGGTGTGAAATACCGCACAGATGCGTAAGGAGAAAATACCGCATCAGGCGCCATTC
GCCATTCAGGCTGCGCAACTGTTGGGAAGGGCGATCGGTGCGGGCCTCTTCGCTATTACGCCAGCTGGCGAAAGGGGGAT
GTGCTGCAAGGCGATTAAGTTGGGTAACGCCAGGGTTTTCCCAGTCACGACGTTGTAAAACGACGGCCAGTGAATTGGAGA
TCGGTACTTCGCGAATGCGTCGAGATTAATACGACTCACTATAGGTTAAAACAGCCTGTGGGTTGTACCCACCCACAGGGCCC
ACTGGGCGCTAGCACACTGGTATCACGGTACCTTTGTGCGCCTGTTTTATACCCCCTTCCCGCAACCGCAAATTTAGAAGCAA
AGCTAACCCGATCGATAGCGGATGCGGCATGCCAGCCGCATTTTGATCAAGTACTTCTGTTTCCCCGGACCGAGTATCAATAG
ACTGCTCACGCGGTTGAAGGAGAAAACGTCCGTTACCCGACCAACTACTTCGAGAAACCTAGTAACATCATGAATGTTGCAG
GGCGTTTCGATCAGCACGACCCTGGTGTAGATCAGGCTGATGAGTCACCGCATTCCCCACGGGTGACCGTGGCGGTGGCTG
CGTTGGCGGCCTGCCTATGGGGTGACCCATAGGACGCTCTAATACGGACATGGTGCGAAGAGTCTATTGAGCTAGTTGGTAG
TCCTCCGGCCCCTGAATGCGGTTAATCCTAACTGCGGAGCACATACCCCTAATCCAAGGGGCAGTGTGTCGTAACGGGCAACT
CTGCAGCGGAACCGACTACTTTGGGTGTCCGTGTTTCCTTTTATTTTTTATACTGGCTGCTTATGGTGACAATCTCAGAGTTGT
TACCATATAGCTATTGGATTGGCCATCCGGTGAGCAACAGAGCTGTCATTTATCAGTTTGTTGGCTTTATACCTCTAAATCACAC
GGTTTTTGGAACGCTTGTATTCATCTTAACCCTCAATAAGGCAAAATGGGAGCGCAAGTATCAACACAAAAGACCGGTGCGC
ATGAAACCGGCTTGAACGCCAGTGGTAGTTCTATAATCCACTACACCAACATAAACTACTATAAAGATGCAGCATCCAACTCGG
CAAATAGGCAAGAATTTTCACAAGACCCTGGTAAGTTCACCGAACCAGTGAAGGATATCATGGTGAAGTCACTACCTGCACTT
AACTCGCCGTCTGCTGAAGAGTGTGGGTACAGTGACAGAGTGCGATCCATAACACTAGGTAACTCTACCATCACAACACAAG
AAAGTGCAAATGTAGTAGTGGGGTATGGTCGATGGCCTGAGTACCTGAAAGACAATGAGGCCACTGCTGAAGATCAACCAA
CCCAGCCTGATGTAGCAACATGTAGGTTTTACACCCTGGAATCGGTCACGTGGGAAAGAGATTCACCCGGGTGGTGGTGGA
AATTCCCGGACGCCCTAAAAGATATGGGGCTCTTCGGCCAAAACATGTACTACCACTATCTAGGAAGAGCCGGTTACACATTG
CATGTACAATGTAATGCATCTAAATTCCATCAGGGATGCTTGCTAGTGGTCTGTGTACCGGAAGCAGAGATGGGATGCAGCCA
AGTGGATCGTACTGTAAATGAGCATGGATTGAGTGAGGGGGAGACCGCTAAGAAATTCTCTTCTACCAGCACAAATGGGACC
AACACGGTACAGACGATTGTGACAAATGCCGGTATGGGAGTGGGAGTGGGCAATCTCACTATATACCCACATCAGTGGATAA
ATTTGCGCACCAATAACTGCGCCACCATCGTCATGCCATACATAAACAACGTACCGATGGACAACATGTTCAGACACCACAATT
TCACACTAATGATTATTCCCTTTGTACCATTAGACTATTCTTCAGATTCATCCACGTACGTGCCCATAACAGTGACAGTCGCTCCA
ATGTGTGCTGAGTATAATGGTTTGAGGCTCTCAACCTCATTGCAAGGATTACCTGTCATGAATACACCGGGTAGCAACCAGTTT
CTGACAACGGACGACTTCCAGTCACCATCTGCCATGCCACAATTTGATGTCACCCCAGAGTTAAATATACCAGGGGAGGTACA
AAACCTCATGGAAATTGCTGAAGTCGACTCGGTGGTGCCAGTCAACAACGTGGAAGGGAAACTCGACACAATGGAGGTCTA
CCGGATTCCAGTGCAGAGTGGTAATCACCAAAGTGACCAAGTCTTCGGTTTTCAAGTGCAACCTGGGCTAGATAGCGTTTTC
AAACACACGCTACTGGGGGAGATTTTGAACTACTATGCACACTGGTCTGGTAGTATAAAACTAACATTTGTTTTCTGTGGTTCC
GCTATGGCTACGGGTAAATTCCTACTAGCCTACGCCCCGCCCGGAGCGAACGCTCCTAAGAATAGGAAAGATGCAATGCTGG
GCACACACATTATCTGGGATGTTGGACTGCAGTCATCGTGTGTCTTATGTGTGCCTTGGATTAGTCAAACTCACTATAGGTTGG
TGCGGCAGGACGAGTACACAAGCGCTGGCAATGTCACATGCTGGTATCAGACTGGAATAGTCGTCCCGGCGGGCACTCCGA
CATCGTGCTCCATCATGTGTTTTGTATCGGCATGCAATGATTTCTCTGTGAGATTACTAAAGGACACGCCATTTATAGAACAAAC
TGCATTACTGCAAGGTGATGTGGTAGAAGCTGTAGAGAACGCCGTTGCACGTGTGGCAGATACAATTGGTAGTGGGCCGTCA
AATTCGCAAGCAGTGCCTGCTTTAACAGCAGTTGAGACAGGGCACACATCTCAGGTGACACCCAGTGATACCATGCAAACCA
GGCATGTCAAGAACTACCATTCCAGATCTGAGTCCAGCATTGAAAACTTCCTCAGCAGATCTGCCTGCGTTTATATGGGAGGA
TACCACACAACCAACACTGACCAGACAAAATTATTTGCCTCATGGACTATTAGTGCACGACGCATGGTTCAAATGAGACGCAA
GCTAGAGATCTTCACTTACGTCCGTTTTGATGTGGAGGTGACTTTTGTGATTACCAGCAAGCAGGACCCGGGCAACCGATTG
GGCCAAGACATGCCACCCCTGACTCACCAGATCATGTATATCCCACCAGGGGGGCCCATTCCAAAGTCTGTCACTGACTATGC
ATGGCAAACCTCCACCAACCCCAGCATTTTCTGGACTGAGGGGAACGCGCCACCCAGAATGTCTATCCCATTCATTAGCATTG
GTAACGCCTACAGTAATTTTTACGACGGGTGGTCTCACTTCTCGCAAAACGGGGTGTATGGCTACAACACACTCAACCACATG
GGTCAAATTTATGTTAGACACGTGAATGGATCATCACCACTCCCTATGACTAGCACTGTTAGAATGTACTTCAAGCCGAAGCAT
GTTAAAGCATGGGTCCCGCGGCCTCCTAGGCTATGCCAATACGAAAATGCATCCACGGTGAACTTTACACCCACAAACGTCAC
CGACAAGCGAACCAGCATCAACTACATTCCTGAGACGGTCAAACCAGACCTATCAAACTACGGAGCTTTTGGACACCAATCA
GGCGCCGTGTACGTCGGAAACTACAGGGTGGTGAACCGGCATCTGGCAACCCATACCGACTGGCAAAACTGTGTATGGGAG
GACTACAACAGAGACCTCCTTATAAGCACCACCACAGCCCACGGATGCGATGTTATAGCCAGGTGTCGCTGTTCAACGGGGG
TCTACTACTGTCAGTCTAAGGGTAAGCACTACCCAGTCAATTTTGAGGGACCGGGTCTTGTGGAAGTTCAGGAGAGTGAGTA
CTATCCCAAGAGATATCAATCCCATGTCCTTCTCGCGGCAGGATTTTCCGAGCCTGGCGATTGTGGTGGGATTCTGAGATGCG
AGCACGGTGTCATCGGAATTGTGACCATGGGGGGTGAAGGTGTCGTTGGCTTTGCCGACGTGCGTGACCTCTTGTGGTTGG
AGGACGATGCGATGGAGCAGGGAGTGAAGGACTATGTAGAGCAACTTGGAAATGCCTTTGGTTCGGGCTTCACCAACCAAA
TTTGTGAACAAGTCAACCTCCTAAAAGAGTCACTAGTGGGTCAAGACTCCATCCTAGAGAAGTCTCTGAAAGCTTTGGTGAA

AATAATATCAGCCTTAGTAATTGTGGTGAGAAATCACGATGACTTAATCACAGTAACTGCCACACTGGCCCTCATCGGCTGCAC
CTCATCCCCGTGGCGGTGGCTTAAACAGAAGGTATCGCAATACTATGGGATACCCATGGCTGAACGTCAAAACAACGGGTGG
CTCAAGAAGTTCACTGAAATGACCAACGCTTGTAAGGGTATGGAATGGATATCCATAAAAATCCAGAAATTCATAGAATGGCT
TAAGGTCAAAATATTACCAGAGGTCAGAGAAAAACATGAATTCCTGAACAGACTCAAGCAGCTCCCTCTGTTGGAAAGCCAG
ATCGCCACAATCGAGCAAAGTGCGCCGTCCCAGAGTGACCAAGAGCAATTGTTTTCCAATGTCCAGTACTTTGCTCACTATTG
CAGGAAGTATGCTCCCCTCTACGCATCAGAAGCAAAGAGAGTATTCTCCCTTGAGAAGAAGATGAGCAATTACATACAGTTCA
AGTCCAAATGCCGTATTGAACCTGTATGTCTACTTCTACACGGGAGCCCTGGCGCCGGTAAGTCGGTAGCAACAAATCTAATC
GGAAGATCACTCGCTGAGAAACTTAACAGCTCAGTGTACTCACTACCACCAGACCCAGATCACTTTGACGGATATAAACAGCA
GGCCGTGGTGATCATGGACGACTTGTGCCAGAATCCTGATGGAAAAGATGTCTCTTTGTTCTGCCAAATGGTCTCTAGTGTAG
ACTTTGTGCCACCTATGGCTGCCTTGGAAGAGAAAGGCATTCTGTTCACTTCTCCATTCGTCCTGGCGTCAACTAACGCAGGG
TCTATCAACGCCCCAACCGTGTCAGATAGTAGGGCCCTGGCGCGAAGGTTCCACTTCGACATGAACATTGAAGTTATCTCCAT
GTACAGCCAAAATGGCAAAATAAATATGCCGATGTCAGTGAAAACGTGTGATGAAGAGTGTTGTCCAGTCAACTTTAAGAGA
TGCTGCCCCCTAGTGTGTGGGAAAGCAATTCAGTTTATAGATAGAAGAACTCAAGTCAGATACTCCCTTGACATGCTGGTAAC
TGAGATGTTCAGGGAATACAATCACAGGCACAGTGTTGGGGCAACCCTTGAAGCGCTATTCCAGGGTCCACCGATATACAGA
GAGATTAAGATCAGCGTGGCACCAGAGACACCACCACCACCAGCCATTGCAGACTTGCTCAAGTCAGTGGACAGTGAAGCC
GTGAGAGAGTACTGTAAAGAAAAGGGGTGGCTGGTTCCAGAGGTTAACTCTACCCTACAGATTGAGAAGTACGTTAGTCGG
GCCTTCATCTGTTTGCAAGCATTGACCACTTTTGTCTCAGTGGCTGGAATTATTTACATAATCTACAAGCTCTTCGCAGGCTTCC
AAGGAGCATACACAGGGATGCCCAATCAGAAACCCAAAGTGCCCACACTCAGGCAGGCAAAAGTGCAAGGACCTGCGTTC
GAATTTGCCGTAGCCATGATGAAGAGGAATTCAAGCACAGTGAAGACCGAATACGGCGAATTCACTATGTTGGGCATTTATG
ACAGGTGGGCCGTACTGCCACGCCATGCTAAACCTGGACCAACCATTCTGATGAACGATCAAGAGGTCGGTGTGCTTGACGC
CAAGGAACTAGTGGATAAGGATGGCACCAATCTGGAGTTGACACTACTCAAGTTAAACCGGAATGAGAAGTTCAGAGACAT
CAGAGGCTTCTTGGCCAAAGAGGAGGTGGAAGCTAACGAGGCTGTACTGGCGATTAACACTAGCAAGTTCCCCAACATGTA
CATCCCAGTGGGTCAAGTCACAGATTACGGTTTCTTAAACTTAGGCGGTACACCCACCAAGAGAATGCTCATGTACAACTTCC
CCACACGAGCGGGCCAGTGCGGCGGGGTTCTCATGTCCACCGGTAAGGTCTTGGGGATCCACGTTGGTGGTAATGGTCATC
AGGGCTTCTCAGCTGCACTCCTCAAGCACTATTTCAATGATGAGCAAGGGGAAATTGAGTTTATTGAGAGTTCAAAGGATGC
GGGGTTCCCAATCATTAATACGCCTAGTAAGACTAAGTTGGAACCGAGCGTCTTCCATCAAGTATTCGAAGGGGATAAAGAAC
CAGCTGTCCTCAGGAACGGTGATCCACGCCTCAAGGCCAACTTTGAGGAAGCCATATTCTCAAAATACATTGGAAATGTCAAC
ACACACGTGGATGAATACATGCTAGAGGCTGTCGATCATTATGCTGGTCAGCTGGCCACACTGGATATCAGCACCGAACCTAT
GAGATTGGAGGATGCTGTGTATGGCACCGAGGGCCTCGAAGCCCTTGACCTAACAACGAGTGCAGGCTACCCTTATGTCGCA
CTAGGCATCAAGAAGAGAGACATCCTTTCAAGGAGGACCAGGGATCTAACCAAGTTGAAGGAATGTATGGATAAATACGGTT
TGAACTTACCAATGGTGACTTATGTGAAAGATGAACTTAGGTCTGCAGACAAAGTAGCAAAAGGGAAGTCTAGGTTGATTGA
AGCATCCAGTTTGAATGACTCTGTAGCAATGAGACAAACATTTGGCAACCTGTACAGAACCTTCCATCTAAACCCAGGGATCG
TGACTGGTAGCGCTGTCGGGTGCGACCCGGACCTCTTTTGGAGTAAAATTCCAGTGATGTTGGATGGTCACCTCATAGCCTTT
GACTACTCTGGATATGATGCTAGCTTGAGCCCCGTGTGGTTTGCCTGCCTAAAACTATTACTTGAGAAATTAGGCTACACACAC
AAGGAAACAAATTACATTGACTACCTGTGTAATTCCCACCACCTGTACAGAGACAAACACTACTTTGTGCGGGGTGGTATGCC
CTCAGGATGTTCCGGCACCAGCATATTTAACTCAATGATAAACAACATCATCATCAGGACTCTCATGCTAAAAGTGTATAAGGG
AATTGATTTGGACCAGTTTAGGATGATTGCATACGGTGACGACGTGATTGCGTCATATCCGTGGCCCATCGATGCATCTTTACT
TGCCGAAGCCGGCAAAGGTTATGGGTTGATTATGACACCAGCAGATAAAGGGGAGTGCTTCAACGAAGTCACCTGGACCAA
CGTCACATTCCTGAAGAGGTATTTTAGAGCAGATGAGCAGTATCCCTTCCTGGTACACCCAGTCATGCCCATGAAAGACATCC
ACGAGTCCATTAGGTGGACCAAAGACCCAAAGAACACCCAAGACCACGTGCGCTCGCTGTGTTTATTGGCCTGGCATAATGG
GGAGCACGAGTATGAGGAGTTCATCCGCAAGATCAGGAGCGTCCCGGTCGGACGTTGCTTGACTCTGCCCGCATTTTCAACC
TTGCGTAGGAAGTGGTTGGACTCTTTTTAAACTAGAGACAATTTGGACTAATTTGAATTGGCTTAACCCTACTGCACTAACCG
AACTAGTCAACGGTGCAGTAGGGGTAAATTCTCCGCATTCGGTGCGAAAAAAAAAAAAAAAAAAAAAAAAAACGCGTATCGG
ATGCCGGGACCGACGAGTGCAGAGGCGTGCAAGCGAGCTTGGCGTAATCATGGTCATAGCTGTTTCCTGTGTGAAATTGTTA
TCCGCTCACAATTCCACACAACATACGAGCCGGAAGCATAAAGTGTAAAGCCTGGGGTGCCTAATGAGTGAGCTAACTCACA
TTAATTGCGTTGCGCTCACTGCCCGCTTTCCAGTCGGGAAACCTGTCGTGCCAGCTGCATTAATGAATCGGCCAACGCGCGG
GGAGAGGCGGTTTGCGTATTGGGCGCTCTTCCGCTTCCTCGCTCACTGACTCGCTGCGCTCGGTCGTTCGGCTGCGGCGAGC
GGTATCAGCTCACTCAAAGGCGGTAATACGGTTATCCACAGAATCAGGGGATAACGCAGGAAAGAACATGTGAGCAAAAGG
CCAGCAAAAGGCCAGGAACCGTAAAAAGGCCGCGTTGCTGGCGTTTTTCCATAGGCTCCGCCCCCCTGACGAGCATCACAA
AAATCGACGCTCAAGTCAGAGGTGGCGAAACCCGACAGGACTATAAAGATACCAGGCGTTTCCCCCTGGAAGCTCCCTCGT
GCGCTCTCCTGTTCCGACCCTGCCGCTTACCGGATACCTGTCCGCCTTTCTCCCTTCGGGAAGCGTGGCGCTTTCTCATAGCTC
ACGCTGTAGGTATCTCAGTTCGGTGTAGGTCGTTCGCTCCAAGCTGGGCTGTGTGCACGAACCCCCCGTTCAGCCCGACCGC
TGCGCCTTATCCGGTAACTATCGTCTTGAGTCCAACCCGGTAAGACACGACTTATCGCCACTGGCAGCAGCCACTGGTAACAG
GATTAGCAGAGCGAGGTATGTAGGCGGTGCTACAGAGTTCTTGAAGTGGTGGCCTAACTACGGCTACACTAGAAGAACAGT
ATTTGGTATCTGCGCTCTGCTGAAGCCAGTTACCTTCGGAAAAAGAGTTGGTAGCTCTTGATCCGGCAAACAAACCACCGCT
GGTAGCGGTGGTTTTTTTGTTTGCAAGCAGCAGATTACGCGCAGAAAAAAAGGATCTCAAGAAGATCCTTTGATCTTTTCTA
CGGGGTCTGACGCTCAGTGGAACGAAAACTCACGTTAAGGGATTTTGGTCATGAGATTATCAAAAAGGATCTTCACCTAGAT

CCTTTTAAATTAAAAATGAAGTTTTAAATCAATCTAAAGTATATATGAGTAAACTTGGTCTGACAGTTACCAATGCTTAATCAGT
GAGGCACCTATCTCAGCGATCTGTCTATTTCGTTCATCCATAGTTGCCTGACTCCCCGTCGTGTAGATAACTACGATACGGGAG
GGCTTACCATCTGGCCCCAGTGCTGCAATGATACCGCGAGACCCACGCTCACCGGCTCCAGATTTATCAGCAATAAACCAGCC
AGCCGGAAGGGCCGAGCGCAGAAGTGGTCCTGCAACTTTATCCGCCTCCATCCAGTCTATTAATTGTTGCCGGGAAGCTAGA
GTAAGTAGTTCGCCAGTTAATAGTTTGCGCAACGTTGTTGCCATTGCTACAGGCATCGTGGTGTCACGCTCGTCGTTTGGTAT
GGCTTCATTCAGCTCCGGTTCCCAACGATCAAGGCGAGTTACATGATCCCCCATGTTGTGCAAAAAAGCGGTTAGCTCCTTCG
GTCCTCCGATCGTTGTCAGAAGTAAGTTGGCCGCAGTGTTATCACTCATGGTTATGGCAGCACTGCATAATTCTCTTACTGTCA
TGCCATCCGTAAGATGCTTTTCTGTGACTGGTGAGTACTCAACCAAGTCATTCTGAGAATAGTGTATGCGGCGACCGAGTTGC
TCTTGCCCGGCGTCAATACGGGATAATACCGCGCCACATAGCAGAACTTTAAAAGTGCTCATCATTGGAAAACGTTCTTCGGG
GCGAAAACTCTCAAGGATCTTACCGCTGTTGAGATCCAGTTCGATGTAACCCACTCGTGCACCCAACTGATCTTCAGCATCTTT
TACTTTCACCAGCGTTTCTGGGTGAGCAAAAACAGGAAGGCAAAATGCCGCAAAAAAGGGAATAAGGGCGACACGGAAAT
GTTGAATACTCATACTCTTCCTTTTTCAATATTATTGAAGCATTTATCAGGGTTATTGTCTCATGAGCGGATACATATTTGAATGTA
TTTAGAAAAATAAACAAATAGGGGTTCCGCGCACATTTCCCCGAAAAGTGCCACCTGACGTCTAAGAAACCATTATTATCATG
ACATTAACCTATAAAAATAGGCGTATCACGAGGCCCTTTCGTC

polyA tail

MluI restriction site

T7 RNA polymerase promotor

Genome of our laboratory E11 strain (GenBank accession number PX993811)

Selected residues for mutagenesis

The E11 genome was mutated as follows:

| Name of Mutant | Position of residue (nt) | Amino acid substitution | Codon in E11 | Codon in Mutant |
|---|---|---|---|---|
| **pE11.VP1.K155N** | 2916-2918 | K → N | AAG | AAC |
| **pE11.VP1.K274E** | 3273-3275 | K → E | AAG | GAA |
| **pE11.VP2.Y97Q** | 1242-1244 | Y → Q | TAC | CAG |
| **pE11.VP3.R69Q** | 1934-1936 | R → Q | CGG | CAG |

### Supplementary Tables

**Supplementary Table S1**. Inactivation rate constants (*k*) of E11, pE11, and E11 mutants in microbially active lakewater. pE11 and E11 mutants were incubated in lakewater collected in September 2025.

| **Sample** | ***k* ± error [h$^{-1}$]** | |
|---|---|---|
| April 2024 | 0.11 | 0.01 |
| July 2024 | 0.09 | 0.01 |
| September 2024 | 0.05 | 0.01 |
| October 2024 | 0.11 | 0.01 |
| February 2025 | 0.09 | 0.01 |
| April 2025 | 0.13 | 0.01 |
| July 2025 | 0.15 | 0.01 |
| September 2025 | 0.19 | 0.01 |
| pE11 | 0.11 | 0.01 |
| pE11.VP1.K155N | 0.18 | 0.02 |
| pE11.VP1.K274E | 0.11 | 0.01 |
| pE11.VP2.Y97Q | 0.01 | 0.01 |
| pE11.VP3.R69Q | 0.11 | 0.01 |

**Supplementary Table S2**. List of all visible P1-P1' (flanking the cleaved peptide bond) and P2-P1 (N-terminal to the cleaved peptide bond) amino acid pairs visible on the proteolytic fingerprints of Lake Geneva. The number of occurrences of each pair in the proteolytic fingerprints is indicated (out of eight lakewater samples). Fully surface-exposed P1 amino acids belonging to a selected P1-P1' or P2-P1 pair are indicated by an asterisk (*). These were selected based on visual inspection of the modeled E11 capsid (**Figure 2A**, **Supplementary Figure 5**). All selected P1 residues (*) were considered potential proteolytic cleavage sites and nominated for mutation. Note that VP3.R69 appears twice as both P2-P1 and P1-P1' pairs surrounding the selected P1 residue are observed in the proteolytic fingerprints.

| P1-P1' pairs | Occurrence (#/8) | Surface-exposed P1 residue (*) | P2-P1 pairs | Occurrence (#/8) | Surface-exposed P1 residue (*) |
|---|---|---|---|---|---|
| A-K | 1 | | M-A | 1 | |
| A-M | 1 | | M-K | 6 | |
| A-R | 1 | | M-R | 6 | |
| K-M | 6 | | M-Y* | 2 | VP2.Y97 |
| K*-R | 8 | VP1.K274 | M-W | 5 | |
| K*-S | 2 | VP1.K155 | Y-A | 1 | |
| K-K | 2 | | Y-K | 2 | |
| K-I | 2 | | Y-R* | 2 | VP3.R69 |
| R-M | 6 | | Y-W | 2 | |
| R-R | 8 | | | | |
| R-S | 2 | | | | |
| R-K | 2 | | | | |
| R*-I | 2 | VP3.R69 | | | |
| W-M | 6 | | | | |
| W-R | 7 | | | | |
| W-S | 2 | | | | |
| W-K | 1 | | | | |
| W-I | 1 | | | | |
| Y-M | 2 | | | | |
| Y-R | 2 | | | | |
| Y-S | 2 | | | | |

**Supplementary Table S3**. List of primers used for E11 full genome sequencing and E3, E7, and CVB5 viral protein sequencing. Primers indicated by an asterisk (*) were solely used for Sanger sequencing and not for genome amplification. The positions of each primer relative to the genome sequence of the corresponding enterovirus is indicated (GenBank accession numbers: X80059 for E11, AY302553 for E3, AF465516 for E7, and MG845891 for CVB5). E: echovirus, CV: coxsackievirus.

| Primer | 5'-Sequence-3' | Orientation | Position |
|---|---|---|---|
| E11_1 | TTAAAACAGCCTGTGGGTTG | Sense | 1-20 |
| E11_2 * | ACTTTGGGTGTCCGTGTTTC | Sense | 549-568 |
| E11_3 * | CGGATGGCCAATCCAATAG | Antisense | 625-643 |
| E11_4 * | TCACGTGGGAAAGAGATTCA | Sense | 1’164-1’183 |
| E11_5 * | TACATGTTTTGGCCGAAGAG | Antisense | 1’229-1’248 |
| E11_6 * | TCATGAATACACCGGGTAGC | Sense | 1’755-1’774 |
| E11_7 * | TTAACTCTGGGGTGACATCAAA | Antisense | 1’826-1’847 |
| E11_8 * | TCGTGCTCCATCATGTGTTT | Sense | 2’363-2’382 |
| E11_9 * | CCTTGCAGTAATGCAGTTTGTT | Antisense | 2’439-2’460 |
| E11_10 | CATTTTCTGGACTGAGGGGA | Sense | 2’965-2’984 |
| E11_11 * | GAGACCACCCGTCGTAAAAA | Antisense | 3’040-3’059 |
| E11_12 * | TAAGGAGGTCTCTGTTGTAG | Antisense | 3’439-3’439 |
| E11_13 | GTGGGCTGTGGTGGTGCTTA | Antisense | 3’459-3’478 |
| E11_14 * | CAGTCAATTTTGAGGGACCG | Sense | 3’552-3’571 |
| E11_15 * | CCAGGCTCGGAAAATCCT | Antisense | 3’640-3’657 |
| E11_16 * | TCATAGAATGGCTTAAGGTCAAAA | Sense | 4’164-4’187 |
| E11_17 * | CTTTCCAACAGAGGGAGCTG | Antisense | 4’235-4’254 |
| E11_18 * | CCCCAACCGTGTCAGATAGT | Sense | 4’767-4’786 |
| E11_19 * | GCCATTTTGGCTGTACATGG | Antisense | 4’836-4’855 |
| E11_20 * | AATCAGAAACCCAAAGTGCC | Sense | 5’357-5’376 |
| E11_21 * | CGGTCTTCACTGTGCTTGAA | Antisense | 5’443-5’462 |
| E11_22 * | TTGAGAGTTCAAAGGATGCG | Sense | 5’967-5’986 |
| E11_23 * | TACTTGATGGAAGACGCTCG | Antisense | 6’030-6’049 |
| E11_24 * | CCAGGGATCGTGACTGGTAG | Sense | 6’554-6’573 |
| E11_25 * | CAAAGGCTATGAGGTGACCA | Antisense | 6’628-6’647 |
| E11_26 * | GTGGACCAAAGACCCAAAGA | Sense | 7’159-7’178 |
| E11_27 * | CTTGCGGATGAACTCCTCAT | Antisense | 7’239-7’258 |
| E11_28 | ATTTACCCCTACTACACCGT | Antisense | 7’402-7’421 |
| E3_1 [9] | CGACAGGATTTCACACAGGA | Sense | 870-890 |
| E3_2 | AGATTGGCATCACACCAGGG | Sense | 1’716-1’735 |
| E3_3 | CAGCAGTAGAGACAGGCCAC | Sense | 2’542-2’561 |
| E3_4 | ATTTCCTCCTGAGCTGCACC | Antisense | 2’747-2’766 |
| E3_5 [9] | GCTTTTCACATACGGGCTAA | Antisense | 3’231-3’211 |
| E3_6 | TGCTGTGGTTGTGCTGACTA | Antisense | 3’445-3’464 |

**Supplementary Table S3**. *Continued*.

| Primer | 5'-Sequence-3' | Orientation | Position |
|---|---|---|---|
| E7_1 | ACGGTACCTTTGTGCGCCT | Sense | 63-81 |
| E7_2 * | TCCAGGCGAAGTACACAACC | Sense | 1'840-1'859 |
| E7_3 * | TGGGACCCATACTCGGATGT | Antisense | 3'177-3'196 |
| E7_4 | CCTGTTGTACACCGGCATCT | Antisense | 3'485-3'504 |
| CVB5_1 | GACCCGGGCAAGTTCACAGA | Sense | 633-652 |
| CVB5_2 | GCTACCAGGTGTCAGCATTGTG | Antisense | 1'487-1'508 |
| CVB5_3 | GCGGAATACAATGGCTTGCGA | Sense | 1'443-1'463 |
| CVB5_4 | GCAATGGCCCTTTCCACTGC | Antisense | 2'208-2'227 |
| CVB5_5 | AGGATGCTCAAAGACACACCCTT | Sense | 2'145-2'167 |
| CVB5_6 | TTACCAATGTACACGGCGCCT | Antisense | 3'062-3'082 |

**Supplementary Table S4**. Primer pairs and cycling parameters used for RT-PCR amplification of E11, E3, E7, and CVB5 segments.

| Virus | Primer pairs | RT step | PCR Amplification | Final Elongation |
|---|---|---|---|---|
| E11 | E11_1E11_13<br>E11_10/E11_28 | 50°C 15 min, 98°C 2min | 30 cycles: 98°C 15 sec, 58°C 15 sec, 72°C 3 min | 72°C 5 min |
| E3 | E3_2/E3_4<br>E3_3/E3_6 | 50°C 15 min, 98°C 2 min | 30 cycles: 98°C 15 sec, 60°C 15 sec, 72°C 1 min | 72°C 5 min |
| E3 | E3_1/E3_5 | 50°C 15 min, 98°C 2 min | 35 cycles: 98°C 30 sec, 58°C 30 sec, 72°C 2 min | 72°C 5 min |
| E7 | E7_1/E7_4 | 50°C 15 min, 98°C 2 min | 35 cycles: 98°C 15 sec, 58°C 15 sec, 72°C 3 min | 72°C 5 min |
| CVB5 | CVB5_1/CVB5_2<br>CVB5_3/CVB5_4<br>CVB5_5/CVB5_6 | 50°C 15 min, 98°C 2 min | 30 cycles: 98°C 15 sec, 60°C 15 sec, 72°C 1 min | 72°C 5 min |

**Supplementary Table S5.** Naturally occurring amino acid substitutions found at the proposed E11 proteolytic cleavage sites in closely-related enterovirus type. E: echovirus, EV-B: enterovirus B, CV: coxsackievirus.

| Selected Residue | Naturally Occurring Substitutions |
|---|---|
| VP1.K155 | N (E3, E7, E12, EV-B101)<br>I (EV-B85)<br>A (CVA9)<br>T (EV-B107)<br>D (E19) |
| VP1.K274 | E (E3, E7, CVB4)<br>S (E12, E19)<br>T (CVA9, EV-B107)<br>V (EV-B101)<br>G(EV-B111) |
| VP2.Y97 | Q (CVA9, CVB1, CVB3, CVB4, CVB5)<br>E (E3)<br>H (E12)<br>M (EV-B85, EV-B107)<br>L (E7)<br>F (E19) |
| VP3.R69 | Q (CVB3, CVB5, EV-B97)<br>K (E12)<br>H (E7) |

#### Supplementary Figures

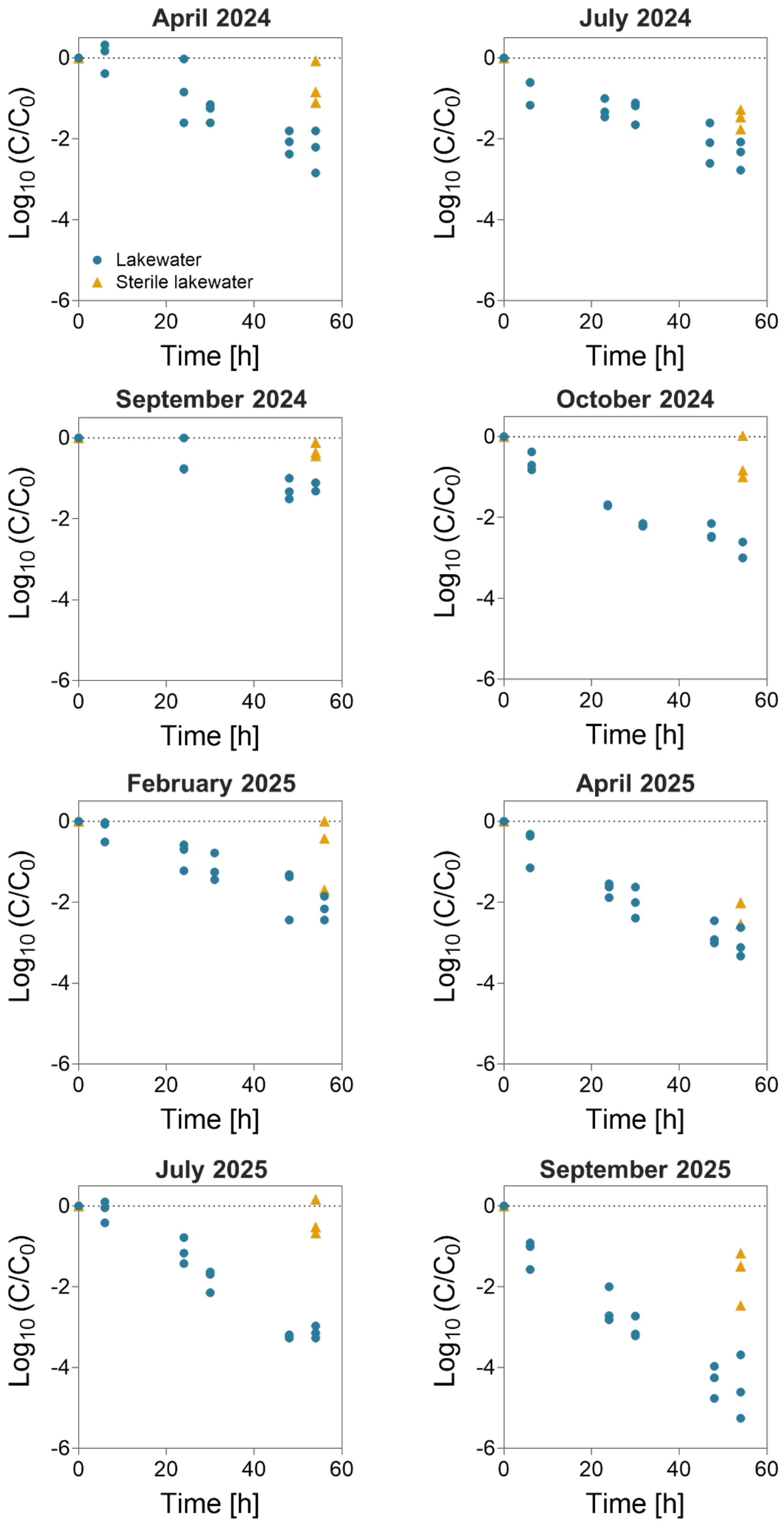


**Supplementary Figure S1**. Echovirus (E11) inactivation in microbially active (circles) and sterile (triangles) Lake Geneva surface water from April 2024 to September 2025. Individual data points of triplicate experiments are presented.

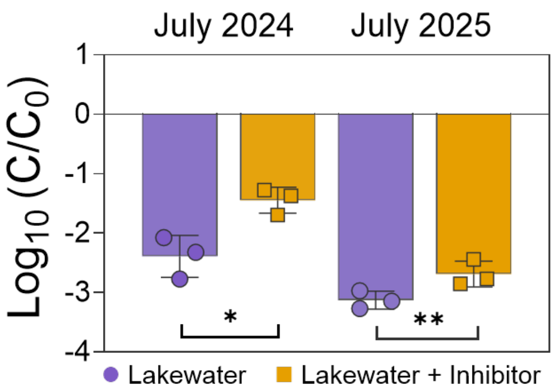


**Supplementary Figure S2**. E11 inactivation in Lake Geneva surface water (t = 54 hours, n = 3) in the absence (circles) or presence (squares) of the metalloprotease inhibitor GM6001. Quantitative results are presented as individual data points with the mean and standard deviation shown by the bar and error bars, respectively. The experiment was performed twice, in July 2024 and July 2025. Significant differences between $\log_{10}$ inactivation were determined using a t-test (*: $p \leq 0.1$; **: $p < 0.05$).

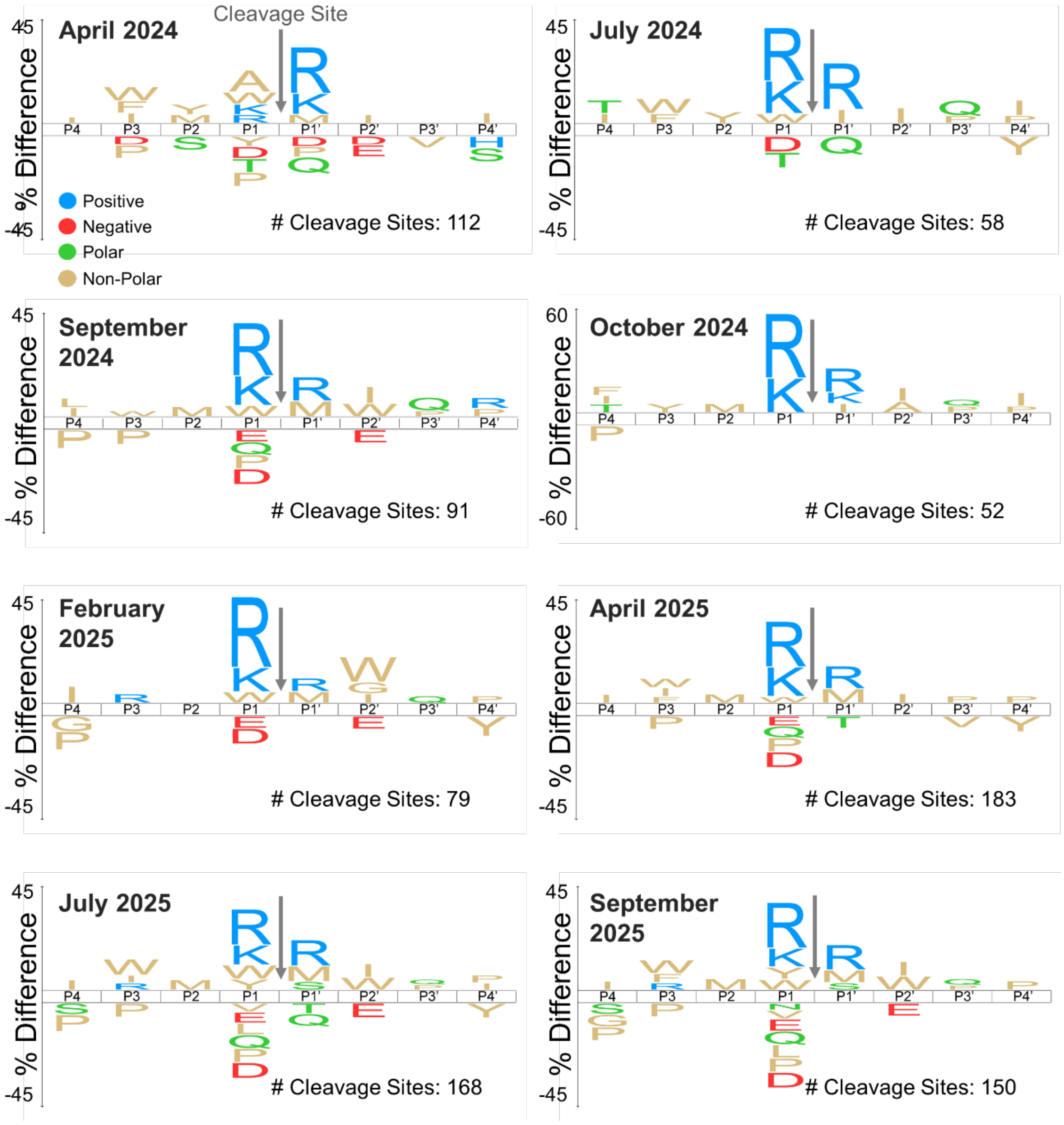


**Supplementary Figure S3**. The proteolytic fingerprints of Lake Geneva surface water from April 2024 to September 2025. April 2024, July 2024, September 2024, October 2024, and February 2025 have been previously reported[10]. The number of proteolytic cleavage sites detected in each sample by the MSP-MS assay, reflective of the heterogeneity of the substrate specificity, is indicated. Note the different y-axis scale for October 2024.

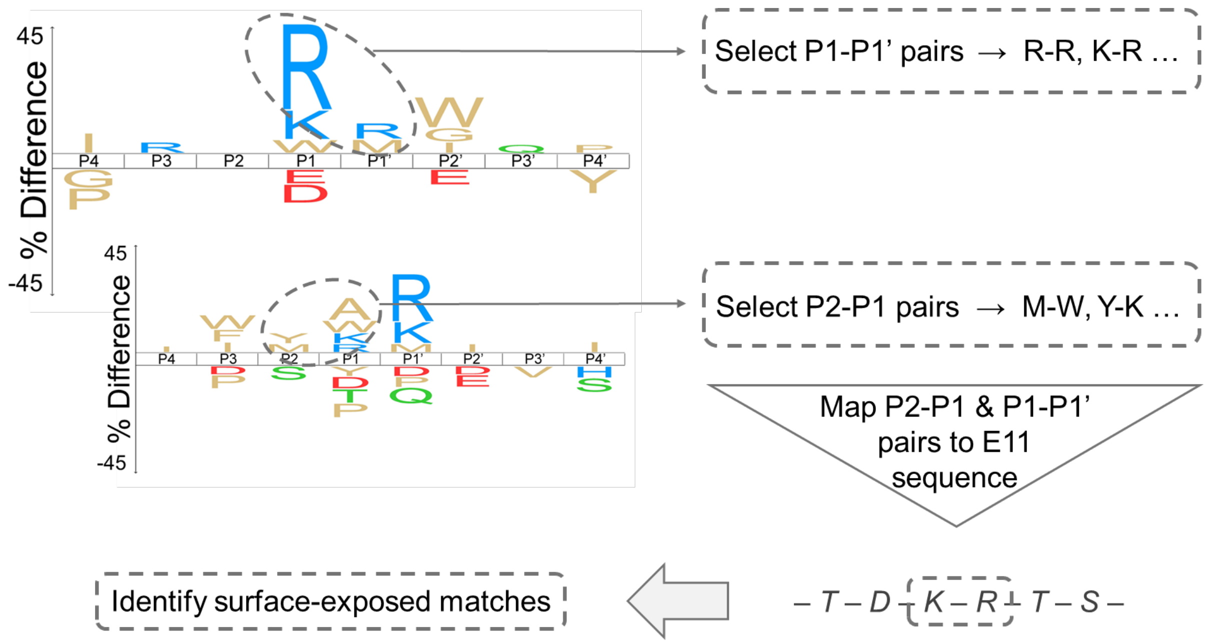


**Supplementary Figure S4**. Scheme of the selection process for potential proteolytic cleavage sites on the surface of E11. Combinations of P1-P1' (flanking the cleaved peptide bond) and P2-P1 (N-terminal to the cleaved peptide bond) amino acid pairs visible on the proteolytic fingerprints of Lake Geneva surface water were identified. These pairs were mapped to the amino acid sequences of the viral proteins (VP1-3). Finally, identification of fully surface-exposed P1 residues of matched pairs led to the selection of potential cleavage sites.

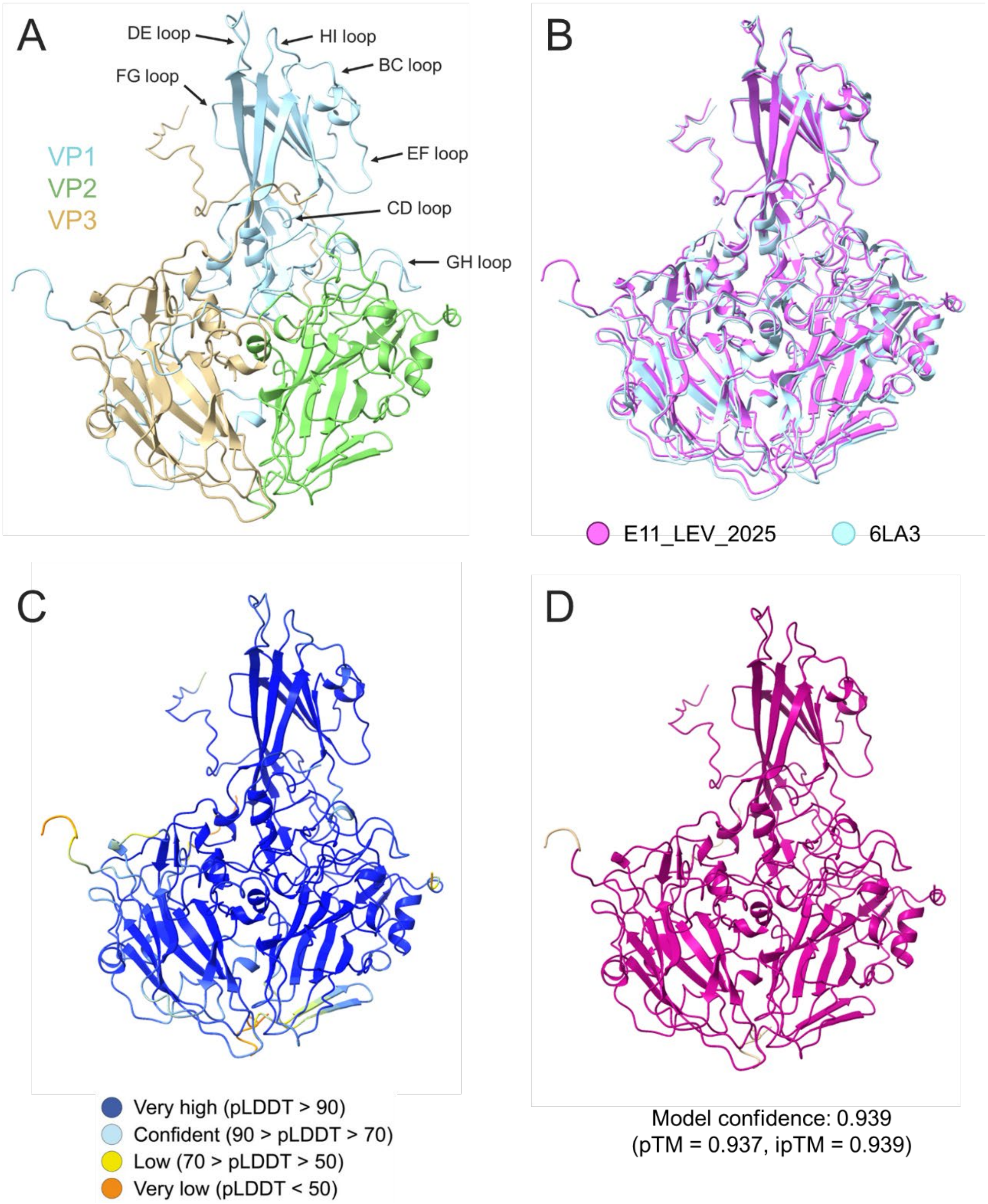


**Supplementary Figure S5**. Ribbon diagram **(A)** of the E11 protomer (rank 1) modeled by AlphaFold2-Multimer displaying the confidence level of the model colored by pLDDT **(C)** or PAE **(D)** score. The model confidence is calculated using Equation S1. Overlaying the predicted E11 protomer (pink) with the structurally determined protomer of E11 (PDB 6LA3, blue) **(B)** revealed a confident match (RMSD = 0.676 Å) between the structures. Images were produced in ChimeraX.

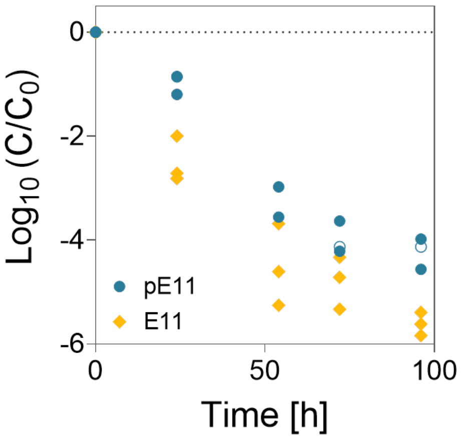


**Supplementary Figure S6**. Inactivation of the plasmid-produced (pE11) and original E11 wild-type virus in Lake Geneva surface water collected in September 2025. Individual data points of triplicate experiments are presented. Both viruses were added to an initial concentration of $10^6$ MPNCU/mL. Values below the detection limit are indicated by open symbols and set to LOQ/√2 where LOQ = 90.4 MPNCU/mL is the limit of detection. No significant difference in inactivation was determined by analysis of covariance (p = 0.279).

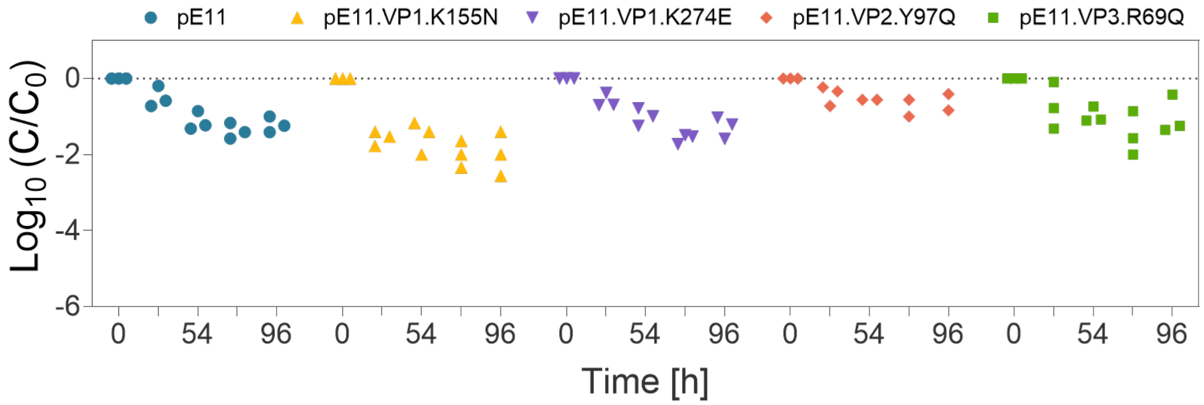


**Supplementary Figure S7**. The inactivation of the plasmid-produced E11 wild-type (pE11) and four mutant viruses in sterile surface water from Lake Geneva. Lakewater was collected in September 2025. All viruses were added to an initial concentration of $10^6$ MPNCU/mL except for pE11.VP2.Y97Q which was added to $10^4$ MPNCU/mL. Individual data points of triplicate experiments are presented.

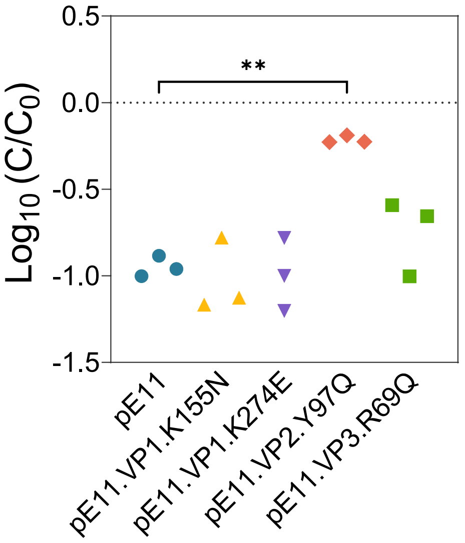


**Supplementary Figure S8**. Inactivation of the plasmid-produced E11 wild-type (pE11) and four mutant viruses in Lake Geneva surface water collected in August 2025 (t = 24 hours). Individual data points of triplicate experiments are presented. Significant differences in $\mathrm{Log}_{10}$ inactivation were determined by one-way analysis of variance with Dunnett's test for multiple comparisons (**: p < 0.01). All viruses were added into lakewater at an initial titer of $10^4$ MPNCU/mL. As in the September 2025 lakewater sample shown in Figure 3 of the main text, the VP2.Y97Q mutation enhances virus stability.

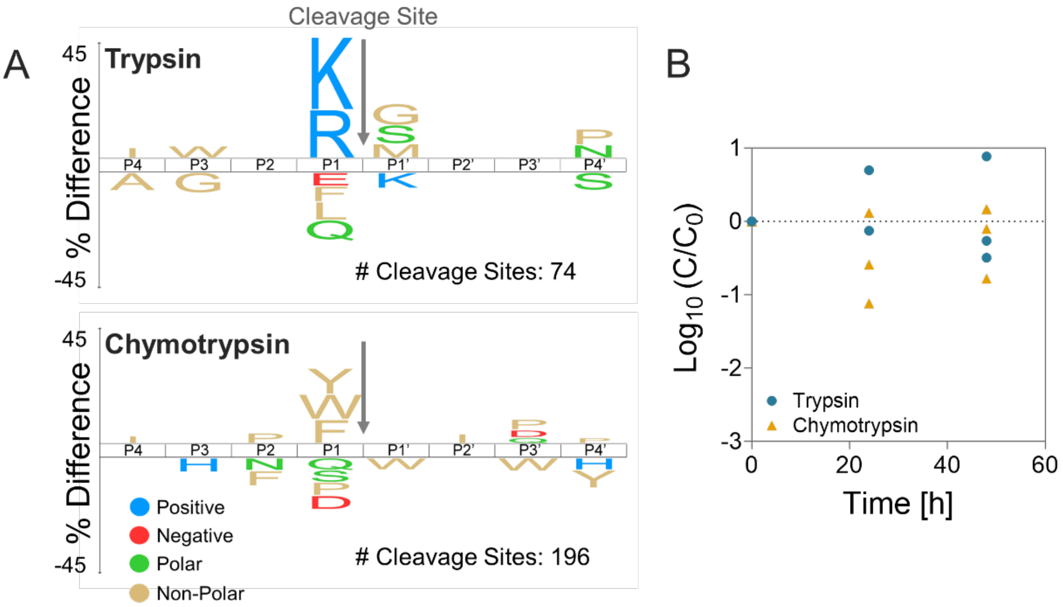


**Supplementary Figure S9**. The effect of two proteases on E11. **(A)** The proteolytic fingerprints of trypsin and chymotrypsin. The number of proteolytic cleavage sites in each sample as detected by the MSP-MS assay is indicated. **(B)** The stability of E11 (t = 48 hours) in the presence of trypsin or chymotrypsin. Individual data points of triplicate experiments are presented.

```
E11        GDVVEAVENAVARVADTIGSGPSNSQAVPALTAVETGHTSQVTPSDTMQTRHVKNYHSRS  60
E3         GDVEEAIDRAVARVADTMPTGPRNTESVPALTAVETGHTSQVVPGDTMQTRHVKNYHSRT  60
E7         GDTDVAINNAVARVADTIASGPSNSTSIPALTAVETGHTSQVEPSDTMQTRHVRNYHSRS  60
CVB5       -PPGEAVERAIARVADTIGSGPVNSESIPALTAAETGHTSQVVPADTMQTRHVKNYHSRS  59
                *::.*:******: :** *: ::*****.******** *.********:*****:

E11        ESSIENFLSRSACVYMGGYHTTNTDQTKLFASWTISARRMVQMRRKLEIFTYVRFDVEVT  120
E3         ESSIENFLCRAACVYITTYKSAGGTPTERYASWRINTRQMVQLRRKFELFTYLRFDMEIT  120
E7         ESTIENFLSRSACVYLEEYFTKDQDDANRYMSWTINARRMVQLRRKFELFTYMRFDMEVT  120
CVB5       ESTVENFLCRSACVFYTTYKNHGT-DGDNFGYWVINTRQVAQLRRKLEMFTYARFDLELT  118
           **::****.*:***:   * . .    . :  * *.:*::.*:***:*:*** ***:*:*
                                             K155
E11        FVITSKQDPGNRLGQDMPPLTHQIMYIPPGGPIPKSVTDYAWQTSTNPSIFWTEGNAPPR  180
E3         FVITSTQDPGTQLAQDMPVLTHQIMYIPPGGPVPNSATDFAWQSSTNPSIFWTEGNAPAR  180
E7         FVITSRQLPGTSIAQDMPPLTHQIMYIPPGGPVPNSVTDFAWQTSTNPSIFWTEGNAPPR  180
CVB5       FVITSTQEQSTIQGQDSPVLTHQIMYVPPGGPVPTKVNSYSWQTSTNPSVFWTEGSAPPR  178
           ***** *  ..  .** * *******:*****:*.....::**:*****:*****.** *

E11        MSIPFISIGNAYSNFYDGWSHFSQNGVYGYNTLNHMGQIYVRHVNGSSPLPMTSTVRMYF  240
E3         MSVPFISIGNAYSNFYDGWSHFTQEGVYGFNSLNNMGHIYVRHVNEQSLGVSTSTLRVYF  240
E7         MSIPFISIGNAYSNFHDGWSHFSQNGVYGYNALNNMGKLYARHVNKDTPYQMSSTIRVYF  240
CVB5       MSIPFISIGNAYSMFYDGWARFDKQGTYGINTLNNMGTLYMRHVNDGSPGPIVSTVRIYF  238
           **:********** *:***::* ::*.** *:**:** :* ****  :     **:*:**
                                            K274
E11        KPKHVKAWVPRPPRLCQYENASTVNFTPTNVTDKRTSINYIPETVKPDLSNY  292
E3         KPKHVRAWVPRPPRLSPYVKSSNVNFKPTAVTTERKDINDVGTLRPMGYTNH  292
E7         KPKHIRVWVPRPPRLCPYIKSSNVNFEPTNLTEKRKSITYVPDTIRPDVRTN  292
CVB5       KPKHVKTWIPRPPRLCQYQKAGNVNFEPTGVTESRTDITTMQ----------  280
           ****::.*:******. * ::..*** ** :* .*..*. :
```

**Supplementary Figure S10**. E11, E3, E7, and CVB5 multiple sequence alignment (Clustal Omega[11]) of viral protein 1 (VP1). The VP1 sequences of our laboratory virus strains were obtained by Sanger sequencing as detailed in the Materials and Methods. Sequence identity with respect to E11 is 71%, 75%, and 69% for E3, E7, and CVB5 respectively. The two hypothesized E11 cleavage sites on VP1 are indicated (boxes).

```
E11          ------SPSAEECGYSDRVRSITLGNSTITTQESANVVVGYGRWPEYLKDNEATAEDQPT  54
E3           -------PTVEECGFSDRVRSITLGNSTITTQECANVVVGYGTWPSYLQDNEATAEDQPT  53
E7 Wallace   -------PSAEECGYSDRVRSLTLGNSTITTQESANVVVGYGRWPEYLRDDEATAEDQPT  53
CVB5         SMPALNSPSAEECGYSDRVRSITLGNSTITTQECANVVVGYGVWPTYLKDDEATAEDQPT  60
                    *:.****:******:***********.******** ** **:*:*********

                                                       Y97
E11          QPDVATCRFYTLESVTWERDSPGWWWKFPDALKDMGLFGQNNYYHYLGRAGYTLHVQCNA  114
E3           QPDVATCRFYTLDSIQWQKESDGWWWKFPEALKNMGLFGQNNEYHYLGRSGYTIHVQCNA  113
E7 Wallace   QPDVATCRFYTLESVQWEKNSAGWWWKFPEALKDMGLFGQNNLYHYLGRAGYTIHVQCNA  113
CVB5         QPDVATCRFYTLESVMWQQSSPGWWWKFPDALSNMGLFGQNNQYHYLGRAGYTVHVQCNA  120
             ************:*: *::.* *******:** .:******** ******:***:******

E11         SKFHQGCLLVVCVPEAEMGCSQVDRTVNEHGLSEGETAKKFSSTSTNGTNTVQTIVTNAG  174
E3          SKFHQGCLLVVCVPEAEMGCSDVEREVVAASLSNEDTAKGFSRTESNGQHTVQTVVYNAG  173
E7 Wallace  SKFHQGCLLVVCVPEAEMGCSQTDKEVAAMNLTKGEAAHKFEPTKTNGEHTVQSIVCNAG  173
CVB5        SKFHQGCLLVVCVPEAEMGCATLANKPDQKSLSNGETANMFESQNSTGQTAVQANVINAG  180
            ********************:    .      .*:: ::*: *.  .:.*  :**: * ***

E11         MGVGVGNLTIYPHQWINLRTNNCATIVMPYINNVPMDNMFRHHNFTLMIIPFVPLDYSSD  234
E3          MGVGVGNLTIFPHQWINLRTNNSATIVMPYINSVPMDNMFRHYNFTLMIIPFAKLEYTEQ  233
E7 Wallace  MGVGVGNLTIYPHQWINLRTNNCATIVMPYVNSVPMDNMFRHYNFTLMVIPFAPLDYAAQ  233
CVB5        MGVGVGNLTIFPHQWINLRTNNSATIVMPYINSVPMDNMFRHNNFTLMIIPFAPLSYSTG  240
            **********:***********.*******:*.********* *****:***. *.*:

E11         SSTYVPITVTVAPMCAEYNGLRLSTSLQ----  262
E3          ASNYVPITVTVAPMCAEYNGLRLASHQ-----  260
E7 Wallace  ASEYVPVTVTIAPMCAEYNGLRLAYQQ-----  260
CVB5        ATTYVPITVTVAPMCAEYNGLRLAGKQGLPTM  272
            :: ***:***:************:
```

**Supplementary Figure S11**. E11, E3, E7, and CVB5 multiple sequence alignment (Clustal Omega[11]) of viral protein 2 (VP2). The VP2 sequences of our laboratory E11, E3, and CVB5 strains were obtained by Sanger sequencing as detailed in the Materials and Methods. Due to difficulty in determining the full VP2 sequence of E7, the VP2 sequence of the E7 reference strain (Wallace, GenBank accession number AY302559) was used. Sequence identity with respect to E11 is 80%, 84%, and 81% for E3, E7, and CVB5 respectively. The hypothesized E11 cleavage site on VP2 is indicated (box).

```
E11      ------------------GLPVMNTPGSNQFLTTDDFQSPSAMPQFDVTPELNIPGEVQN  42
E3       ------------------GLPTMLTPGSNQFLTSDDFQSPSAMPQFDVTPEMKIPGEVHN  42
E7       ------------------GLPVLNTPGSNQFMTSDDFQSPSAMPQFDVTPHMDIPGEVHN  42
CVB5     TVAPMFAEYNGLRLAGKQGLPTMLTPGSNQFLTSDDFQSPSAMPQFDVTPEMDIPGQVNN  60
                           ***.: *******:*:****************.:.***:*:*
                                   R69
E11      LMEIAEVDSVVPVNNVEGKLDTMEVYRIPVQSGNHQSDQVFGFQVQPGLDSVFKHTLLGE  102
E3       LMEIAEVDSVVPVNNTKENVNSMEAYRIPVTGGDQLHTQVFGFQMQPGLNSVFKRTLLGE  102
E7       LMEIAEVDSVVPVNNTQNNLQSMDAYHIEVNTGNHQGDKIFAFQIQPGLDSVFKRTLLGE  102
CVB5     LMEIAEVDSVVPVNNTEGKVLSIESYQIPVQSNSTNGSQVFGFPLTPGASSVLNRTLLGE  120
         ***************.: :: ::: *:* *  ..    ::*.* : ** .**:::*****

E11      ILNYYAHWSGSIKLTFVFCGSAMATGKFLLAYAPPGANAPKNRKDAMLGTHIIWDVGLQS  162
E3       ILNYYAHWSGSVKLTFVFCGSAMATGKFLLAYSPPGASPPQNRKQAMLGTHVIWDVGLQS  162
E7       VLNYYAHWSGSIKLTFTFCGSAMATGKLLLAYSPPGADVPASRKQAMLGTHIIWDIGLQS  162
CVB5     ILNYYTHWSGSIKLTFMFCGSAMATGKFLLAYSPPGAGTPTTRKEAMLGTHVIWDVGLQS  180
         :****:*****:**** **********:****:****. * .**:******:***:****

E11      SCVLCVPWISQTHYRLVRQDEYTSAGNVTCWYQTGIVVPAGTPTSCSIMCFVSACNDFSV  222
E3       SCVLCIPWISQTHYRLVQQDEYTSAGYVTCWYQTGLIVPPGAPPSCTILCFASACNDFSV  222
E7       SCVLCIPWISQTHYRLVQQDEYTSAGNVTCWYQTGIVVPPGTPNKCVVLCFVSACNDFSV  222
CVB5     SCVLCIPWISQTHYRYVVMDEYTAGGYITCWYQTNIVVPADTQSDCKILCFVSACNDFSV  240
         *****:********* *  ****:.* :******.::** .:  .* ::**.********

E11      RLLKDTPFIEQTALLQ  238
E3       RMLRDTPFIEQTQLLQ  238
E7       RMLRDTPFIGQTALLQ  238
CVB5     RMLKDTPFIKQ-----  251
         *:*:***** *
```

**Supplementary Figure S12**. E11, E3, E7, and CVB5 multiple sequence alignment (Clustal Omega[11]) of viral protein 3 (VP3). The VP3 sequences of our laboratory virus strains were obtained by Sanger sequencing as detailed in the Materials and Methods. Sequence identity with respect to E11 is 81%, 81%, and 77% for E3, E7, and CVB5 respectively. The hypothesized E11 cleavage site on VP3 is indicated (box).

#### References


1. Jumper, J. *et al.* Highly accurate protein structure prediction with AlphaFold. *Nature* **596**, 583–583 (2021).
2. Evans, R. *et al.* Protein complex prediction with AlphaFold-Multimer. 2021.10.04.463034 Preprint at https://doi.org/10.1101/2021.10.04.463034 (2022).
3. Mirdita, M. *et al.* ColabFold: making protein folding accessible to all. *Nature Methods* **19**, 679–682 (2022).
4. Mirdita, M., Steinegger, M. & Söding, J. MMseqs2 desktop and local web server app for fast, interactive sequence searches. *Bioinformatics* **35**, 2856–2858 (2019).
5. The UniProt Consortium. UniProt: the universal protein knowledgebase in 2021. *Nucleic Acids Res* **49**, 480–489 (2021).
6. Berman, H. M. *et al.* The Protein Data Bank. *Nucleic Acids Res* **28**, 235–242 (2000).
7. Pettersen, E. F. *et al.* UCSF ChimeraX: Structure visualization for researchers, educators, and developers. *Protein Sci* **30**, 70–82 (2021).
8. Niu, S. *et al.* Molecular and structural basis of Echovirus 11 infection by using the dual-receptor system of CD55 and FcRn. *Chinese Science Bulletin* **65**, 67–79 (2020).
9. Kyriakopoulou, Z. *et al.* Full-Genome Sequence Analysis of a Multirecombinant Echovirus 3 Strain Isolated from Sewage in Greece. *Journal of Clinical Microbiology* **48**, 1513–1519 (2010).
10. Meibom, J., Wichmann, N., Astorch-Cardona, A., Zumstein, M. & Kohn, T. Proteolytic Activity and Substrate Specificity of Lake Geneva. *Environ. Sci. Technol.* **59**, 27811–27823 (2025).
11. Madeira, F. *et al.* The EMBL-EBI Job Dispatcher sequence analysis tools framework in 2024. *Nucleic Acids Res* **52**, 521–525 (2024).
